# Acetylation-Dependent Recruitment of the FACT Complex and Its Role in Regulating Pol II Occupancy Genome-Wide in *Saccharomyces cerevisiae*

**DOI:** 10.1101/285932

**Authors:** Rakesh Pathak, Priyanka Singh, Sudha Ananthakrishnan, Sarah Adamczyk, Olivia Schimmel, Chhabi K. Govind

## Abstract

Histone chaperones, chromatin remodelers, and histone modifying complexes play a critical role in alleviating the nucleosomal barrier for DNA-dependent processes. Here, we have examined the role of two highly conserved yeast (*Saccharomyces cerevisiae*) histone chaperones, FACT and Spt6, in regulating transcription. We show that the H3 tail contributes to the recruitment of FACT to coding sequences in a manner dependent on acetylation. We found that deleting a H3 HAT Gcn5 or mutating lysines on the H3 tail impairs FACT recruitment at *ADH1* and *ARG1* genes. However, deleting the H4 tail or mutating the H4 lysines failed to dampen FACT occupancy in coding regions. Additionally, we show that FACT-depletion reduces Pol II occupancy in the 5’ ends genome-wide. In contrast, Spt6-depletion leads to reduction in Pol II occupancy towards the 3’ end, in a manner dependent on the gene-length. Severe transcription and histone eviction defects were also observed in a strain that was impaired for Spt6 recruitment (*spt6*Δ202) and depleted of FACT. Importantly, the severity of the defect strongly correlated with WT Pol II occupancies at these genes, indicating critical roles of Spt6 and Spt16 in promoting high-level transcription. Collectively, our results show that both FACT and Spt6 are important for transcription globally and may participate during different stages of transcription.

## INTRODUCTION

The nucleosome is the fundamental unit of chromatin and is composed of ∼ 147 bp of DNA wrapped around a histone octamer consisting of two copies of histones H2A, H2B, H3 and H4. Nucleosomes pose a significant impediment to all steps of transcription, including the steps of initiation and elongation of RNA polymerase II (Pol II) through the coding sequences (CDS) (LI *et al*. 2007). There are two principle mechanisms suggested to alleviate nucleosomal barrier: I) removal of the H2A-H2B dimer to generate hexamers that can be readily overcome by elongating polymerases (KIREEVA *et al*. 2002), and II) complete removal of histone octamers leading to reduced nucleosomal density across the transcribing genes (LEE *et al*. 2004; DION *et al*. 2007).

Various factors and enzyme complexes have been implicated in removing the nucleosomal barriers *in vivo* (LI *et al*. 2007). For example, acetylation of histone tails facilitates histone eviction by weakening histone-DNA interactions (GOVIND *et al*. 2007; GOVIND *et al*. 2010) and by promoting recruitment of ATP-dependent remodelers, such as RSC and SWI/SNF (HASSAN *et al*. 2001; DECHASSA *et al*. 2010; SPAIN *et al*. 2014). While nucleosome disassembly is important for transcription, it is also important to restore the chromatin structure in the wake of transcription. Histone chaperones, many of which are cotranscriptionally recruited to actively transcribing genes, are implicated in performing this function (GURARD-LEVIN *et al*. 2014).

Two such chaperones, the FACT complex (Spt16/SSRP1 in humans and Spt16/Pob3 in yeast) and Spt6, are enriched in transcribed coding regions (ANDRULIS *et al*. 2000; KAPLAN *et al*. 2000; KROGAN *et al*. 2002; MASON AND STRUHL 2003; MAYER *et al*. 2010; FORMOSA 2012; BURUGULA *et al*. 2014), consistent with a role for these factors in regulating transcription and in maintaining chromatin integrity. Human FACT recognizes and displaces one of the H2A/H2B dimers from the nucleosome, and promotes transcription on a chromatin template, *in vitro*. In addition, it can assemble all four histones on the DNA (BELOTSERKOVSKAYA *et al*. 2003). Yeast (*S. cerevisiae*) FACT also interacts with all four histones and displays a strong affinity toward intact nucleosomes (FORMOSA *et al*. 2001; VANDEMARK *et al*. 2008). Multiple domains within the FACT subunits Spt16 and Pob3 are implicated in binding and chaperoning histones (HONDELE AND LADURNER 2011; FORMOSA 2012). The N-terminal domain (peptidase-like) of *S. pombe* Spt16, for example, binds to both core histones as well as histone N-terminal tails (STUWE *et al*. 2008), and the M-domain of *Chaetomium thermophilum* Spt16 recognized H2A/H2B with affinity similar to that observed with full-length Spt16 (HONDELE *et al*. 2013). However, in *S. cerevisiae*, the C-terminal regions of both Spt16 and Pob3 were defined as H2A/H2B binding domains (VANDEMARK *et al*. 2008; KEMBLE *et al*. 2015). Whether these different domains are required for interacting with nucleosomes in a context-dependent manner remains to be seen.

FACT is shown to promote reassembly of the displaced H3/H4 in the *ADH1*, *ADH2* and *STE3* coding regions (JAMAI *et al*. 2009) implicating FACT in restoring chromatin behind elongating Pol II. The role of FACT (and Spt6) in regulating histone occupancy has been examined at a genome-wide scale (VAN BAKEL *et al*. 2013; JERONIMO *et al*. 2015). Impairing FACT function leads to reduced histone occupancy and also results in aberrant incorporation of a histone variant H2AZ (JERONIMO *et al*. 2015). Additionally, recruitment of FACT to the *HO* promoter is shown to promote histone eviction (TAKAHATA *et al*. 2009). Likewise, FACT helps in evicting H2A/H2B from the promoter of the *PHO5* gene, upon induction (RANSOM *et al*. 2009).

Gene-specific studies have shown a role for FACT in regulating transcription. For example, FACT mutants impaired Pol II and TBP recruitment at the *GAL1* promoter, implicating a role for FACT in regulating transcription at the initiation step (BISWAS *et al*. 2006; FLEMING *et al*. 2008). However, moderate reductions in Pol II occupancy were observed in the 3’ ORFs of *GAL1* and *PHO5* but not in *LacZ* or *YAT1* genes expressed on plasmids under the control of *GAL1* promoter (JIMENO-GONZALEZ *et al*. 2006). Likewise, transcription defects were observed only at *ADH1* and not at *ADH2* or *STE3*, despite all three genes showing a histone reassembly defect (JAMAI *et al*. 2009), making it unclear if FACT is generally required for transcription. In addition to transcription initiation, FACT is shown to participate in promoting Pol II processivity and elongation rate at *GAL1* gene, in conjunction with the H2B ubiquitination (FLEMING *et al*. 2008).

FACT shares many functional similarities with another histone chaperone, known as Spt6, which interacts with H3/H4 (KAPLAN *et al*. 2003; MAYER *et al*. 2010), and also with H2A/H2B (MCCULLOUGH *et al*. 2015). Loss of Spt6 function leads to reduced histone occupancy over transcribed regions, suggesting a role for Spt6 in cotranscriptional histone reassembly (IVANOVSKA *et al*. 2011; PERALES *et al*. 2013; VAN BAKEL *et al*. 2013; JERONIMO *et al*. 2015). Spt6, along with the FACT complex, is also implicated in preventing spurious incorporation of H2AZ in coding regions (JERONIMO *et al*. 2015) and thereby helps in restricting H2AZ to promoter nucleosomes (BILLON AND COTE 2013). Therefore, Spt6 and FACT play important roles in maintaining chromatin integrity. Consistent with this, widespread aberrant transcription was observed in the cells deficient of Spt6 or FACT (CHEUNG *et al*. 2008; VAN BAKEL *et al*. 2013). Altered histone modifications, and reduced transcription genome-wide has been observed in Spt6 mutant (DEGENNARO *et al*. 2013; KATO *et al*. 2013; PERALES *et al*. 2013). Spt6 possesses a tandem SH2 (tSH2) domain at its C-terminus that interacts with phosphorylated Pol II CTD, *in vitro* (DENGL *et al*. 2009; CLOSE *et al*. 2011; LIU *et al*. 2011) and this domain is implicated in promoting Spt6 recruitment to transcribed genes, *in vivo* (MAYER *et al*. 2010; MAYER *et al*. 2012; BURUGULA *et al*. 2014).

In this study, we show that the acetylated histone H3 tail contributes to efficient recruitment of FACT to transcribed coding sequences in *S. cerevisiae*. Depleting Spt16 subunit of FACT elicits a greater reduction in Pol II occupancies in the 5’ends of the genes, suggesting a potential role for FACT in the early elongation steps of transcription. In contrast, Pol II occupancies were reduced more towards the 3’ end in Spt6-depleted cells, suggestive of processivity defects. Significant reductions in Pol II occupancies were also observed in a strain defective for Spt6 recruitment. Our study reveals that both FACT and Spt6 promotes transcription genome-wide and perhaps participate in different stages of transcription.

## MATERIALS AND METHODS

### Yeast strains and growth conditions

The yeast strains used in this study are listed in Table S1. The cells were grown to absorbance A_600_ of 0.5-0.6 in synthetic complete (SC) media lacking isoleucine/valine, and treated with sulfometuron methyl (SM; 0.6 µg/ml) for 30 minutes to induce Gcn4 targets. Esa1 in the *gcn5*Δ/*esa1ts* strain was inactivated by shifting the cultures grown at 25°C to 37°C for ∼1.5 hours prior to induction by SM. Spt16 and Spt6 were depleted in *SPT16-TET* and *SPT6-TET* (HUGHES *et al*. 2000; MNAIMNEH *et al*. 2004) cells by growing these strains in the presence of 10 μg/ml doxycycline overnight, and sub-culturing the overnight cultures in 100 ml of synthetic complete (SC) media with 10 μg/ml doxycycline to an OD_600_ of 0.6.

### Coimmunoprecipitation Assay

The coimmunoprecipitation experiments were performed as described previously (GOVIND *et al*. 2010). The HA-tagged H2B or Spt16-Myc tagged WT and *H3*Δ*1-28* strains were resuspended in 500 µl of lysis buffer (50 mM Tris-HCl [pH-7.5], 50 mM HEPES-KOH [pH 7.9], 10 mM MgSO_4_, 100 mM (NH_4_)_2_SO_4_, 12.5 mM KOAc, 0.01% NP-40, 20% Glycerol, 1ug/ml Pepstatin A, 100 mM PMSF, 1ug/ml Leupeptin;) and 500 µl of glass beads, and disrupted by vortexing (18 seconds × 8 times, and 150 seconds on ice between each agitation cycle). Whole cell extracts were incubated overnight with magnetic beads that were pre-conjugated to anti-Myc or anti-HA antibodies in 100 µl 4X MTB buffer (200 mM HEPES-KOH [pH 7.9], 800 mM KOAc, 54 mM MgOAc2, 40% Glycerol, 0.04% NP-40, 400 mM PMSF, 4 ug/ml Pepstatin, 4ug/ml Leupeptin), and washed 5 times with the wash buffer (50 mM Tris-HCl (pH-8.0), 0.3% NP-40, 500 mM NaCl, 10% Glycerol, 1mM PMSF, 1ug/ml Leupeptin, and 1ug/ml Pepstatin). Immunoprecipitates were analyzed by western blot using the following antibodies: anti-Myc (Roche), anti-HA (Roche), anti-Spt16 and anti-Spt6 antibodies (kindly provided by Tim Formosa). The signal intensities were quantified using Image Studio lite version 5.2 (LI-COR Biosciences).

### ChIP and ChIP-chip

The cultures were crosslinked with formaldehyde and processed for chromatin immunoprecipitation as described previously (GOVIND *et al*. 2012). ChIPs were performed using antibodies, anti-Myc (Roche), anti-Rpb3 (Neoclone), and anti-H3 (Abcam). ChIP DNA and the related input DNA was amplified using the primers against specific regions. 5µl of ChIP dye (15% Ficol, 0.25% bromophenol blue in 1X TBE) and SYBR green dye were added in the PCR products, resolved on 8% TBE gels and visualized on a phosphorimager and quantified using ImageQuant 5.1 software. The fold enrichments were determined by taking the ratios of the ChIP signal for gene of interest, and the signal obtained for the *POL1* used as an internal control and dividing by the ratios obtained for the related input samples (ChIP/Input fold enrichment: ChIP intensities [*ARG1/POL1*]) / input intensities [*ARG1/POL1*]). The ChIP experiments were performed using at least three independent cultures, and PCR reactions were conducted at least in duplicates. The error bars represent standard error of mean (SEM).

For ChIP-chip experiments, ChIP and related input DNA samples were amplified, from at least two biological replicates, using the GenomePlex complete whole genome amplification (WGA) kit (Sigma, cat # WGA2), according to the manufacturer’s instructions. The amplified ChIP DNA and input DNA were purified by using PCR cleanup kit (Qiagen Cat # 28104) and the DNA was quantified by NanoDrop. The samples were hybridized on Agilent 4×44 arrays (G4493A) after labeling the ChIP and input DNA with Alexa555 and Alexa647 fluorescent dyes, respectively, as per the manufacturer’s instructions at genomic core facility at Michigan State University. The arrays were scanned using Agilent scanner, and data was extracted with the Feature Extraction software (Agilent) as described previously (SPAIN *et al*. 2014).

### Bioinformatics Analysis

The data extracted with the Feature Extraction software (Agilent) was normalized using Limma package from Bioconductor, as described previously (VENKATESH *et al*. 2012). The genes were divided into 10 equal sized bins, with the two bins assigned to the region 500 bp upstream of the transcription start site (TSS) and two bins to the 500 bp to the region downstream of the transcription end site (TES). The average probe enrichment values were assigned to the closest bin according to the probe location, and a 10 bin matrix was generated using a PERL script. Genes corresponding to the majority of dubious ORFs, tRNA genes, small nuclear RNA genes as well as autonomously replicating sequences (ARS) were removed from the dataset. The enrichments in the 6 bins between TSS and TES were averaged to obtain an average ORF occupancy. The genes for analysis were selected on the basis of ORF enrichment. The genes <500 bp in length were removed from the analyses. The versatile aggregate profiler (BRUNELLE *et al*. 2015) was used to generate gene-average profile. The genes were split in the middle, and the probe intensities were aligned to the TSS for the first half and to the TES for the second half of the genes.

### Box-plots

Center lines show the median and box limits indicate the 25th and 75th percentiles as determined by R software. Whiskers extend 1.5 times the interquartile range from the 25th and 75th percentiles and outliers are represented by dots.

### Data Availability

Strains generated in this study, and the sequences of primers used for ChIP analysis are available upon request. The Gene Expression Omnibus accession number for the ChIP-chip data reported in this paper is GSE69642.

## RESULTS

### Histone H3 N-terminal tails facilitate FACT interaction with chromatin *in vivo*

While Spt16 is enriched in coding regions of actively transcribing genes (MASON AND STRUHL 2003; MAYER *et al*. 2010), the mechanisms by which it is recruited remains to be established. Given that the FACT complex interacts with nucleosomes *in vitro* (BELOTSERKOVSKAYA *et al*. 2003), it can be recruited through nucleosome interactions. Moreover, Spt16 interaction was greatly reduced with the nucleosomes lacking the histone N-terminal tails (NTTs) implicating histone tails in promoting Spt16-nucleosome interactions (STUWE *et al*. 2008; VANDEMARK *et al*. 2008). Significantly, removal of the H3 and H4 tails, but not of the H2A-H2B tails, abolished Spt16-histone interactions *in vitro* (WINKLER *et al*. 2011). The H3 or H4 tails therefore could facilitate recruitment/retention of the FACT complex. To test this possibility, we examined yeast Spt16 occupancy by ChIP in histone mutants lacking the H3 or H4 N-terminal tail. Spt16 occupancy in the *ARG1* 5’ and 3’ ORFs was reduced by ∼50% in the H3 mutants lacking 1-20 (*H3Δ1-20*) or 1-28 (*H3Δ1-28*) N-terminal residues (Figure 1A). In comparison, in the H4 tail mutant (*H4Δ1-16*), only a small (∼20%) reduction was observed in the *ARG1* 3’ ORF (Figure 1A). A substantial reduction in Spt16 occupancy (∼80%) was also observed at a constitutively expressed *ADH1* gene in the H3 mutants (Figure 1B), but not in the H4 mutant (Figure S1A). Although, FACT/Spt16 interacts with both H3 and H4 tail peptides, *in vitro*, (STUWE *et al*. 2008; VANDEMARK *et al*. 2008) it appears that the H3 tail may help in recruiting or retaining FACT to its target genes, *in vivo*. Consistent with this idea, reduced occupancy of Spt16 was also observed in the coding regions of *PYK1*, *PMA1* and *GLY1* genes (Figure 1C). In contrast to the impaired Spt16 occupancy in the H3 mutant, for most genes, Pol II occupancies were comparable in WT and the H3 mutants, except for the *GLY1* gene, which showed a small reduction in Pol II occupancy (Figure S1B).

**Figure 1:**
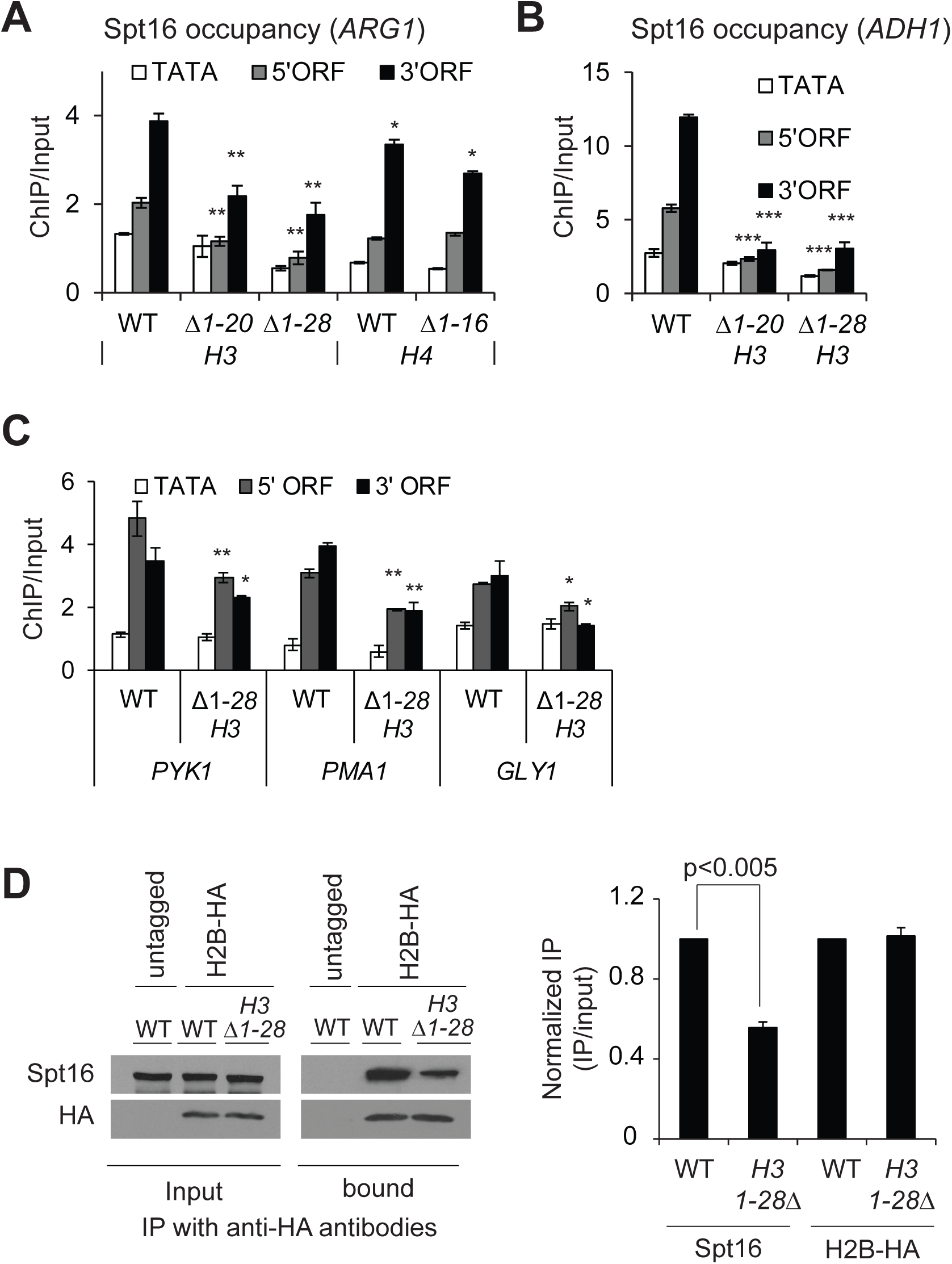
The H3 N-terminal tail promotes FACT recruitment. A-B) ChIP occupancy of Myc-tagged Spt16 in WT and mutants lacking the H3 N-terminal tail residues 1-20 (*H3Δ1-20*) or 1-28 (*H3Δ1-28*) and 1-16 residues of H4 (*H4Δ1-16*) at *ARG1* (A) and in H3 mutants at *ADH1* (B). Graphs show mean and SEM. * represents a p-value < 0.01, ** represents a p-value < 0.001, and *** represents a p-vlaue <0.0001. C) ChIP occupancies of Myc-tagged Spt16 at the indicated genes in the WT and the H3 tail mutant, *H3*Δ*1-28*. * represents a p-value < 0.01, ** represents a p-value < 0.001. D) Whole-cell extracts prepared from HA-tagged H2B WT and *H3Δ1-28* strains were pulled-down with the anti-HA beads, and the immunoprecipitates were subjected to western blot using antibodies against Spt16 and HA. Untagged WT was used as a control. The representative blot is shown on the left, and the quantified data on the right.

To further examine the role of the H3 tails in promoting FACT association with chromatin, we performed coimmunoprecipitation assay. The HA-tagged histone H2B (H2B-HA) efficiently pulled-down Spt16 from the whole cell extracts (WCEs) prepared from HA-tagged WT cells but not from untagged cells (Figure 1D, left). We also observed a reduced Spt16 pull-down from the *H3*Δ*1-28* WCEs (∼50 %; Figure 1D, right). No such reduction in Spt16 occupancy was seen in the H4 tail deletion mutant (*H4*Δ*1-16*; Figure S1C), supporting the idea that the H3 tail promotes Spt16 association with chromatin *in vivo*. However, the extent to which the H3 tail contributes in this process may be variable, as observed by the differences of Spt16 occupancies at the different genes in the H3 mutant (Figure 1A-1C).

### Acetylation of the H3 tail promotes FACT occupancy at *ADH1* and *ARG1* genes

After observing that the H3 tail contributes to Spt16, we examined the role for posttranslational modifications on the H3 tail in regulating FACT localization to transcribed genes. The H3 tail lysines, K4 and K36, are methylated by Set1 and Set2, respectively (BRIGGS *et al*. 2001; STRAHL *et al*. 2002). To examine, whether the reduction in Spt16 occupancy in the H3 tail deletion mutant is due to the loss of H3 methylation, we measured Spt16 enrichment in the *set1*Δ/*set2*Δ double mutant. Comparable occupancies of Spt16 were observed in the ORFs of *ARG1* and *ADH1* genes in WT and the *set1*Δ/*set2*Δ mutant (Figure 2A). Furthermore, we found very similar signal for coimmunoprecipitation of Pol II with Spt16 in WT and *set1*Δ/*set2*Δ WCEs (Figure 2B), suggesting that H3 methylation is likely dispensable for maintaining WT level of FACT occupancy, at least, at these two genes. The H3 and H4 tails are also acetylated by Gcn5-containing SAGA and Esa1-containing NuA4 histone acetyltransferase (HAT) complexes, respectively. Accordingly, a *gcn5*Δ/*esa1ts* double mutant elicits strong reductions in H3 and H4 acetylation (GINSBURG *et al*. 2009). The *gcn5*Δ/*esa1ts* mutant, grown at the permissive temperature 25°C (represented as *gcn5*Δ) or at the non-permissive temperature 37°C to inactivate Esa1 (*gcn5*Δ/*esa1ts*), produced comparable reductions in Spt16 occupancy in the *ARG1* and *ADH1* ORFs (Figure 2C and Figure S2, right). In contrast, both WT and the HAT mutant displayed comparable Pol II occupancies at *ARG1* and *ADH1* (Figure 2D, and Figure S2, left). To examine whether FACT occupancy correlates to H3 acetylation levels, we determined Spt16 enrichment at *ARG1* in a histone deacetylase mutant, *rpd3*Δ/*hos2*Δ. Interestingly, Spt16 occupancy was not increased in the histone deacetylase mutant *rpd3*Δ/*hos2*Δ (Figure 2E). This is surprising given that H3 acetylation levels were shown to be elevated in this mutant (GOVIND *et al*. 2010), and that our current results reveal diminished FACT occupancy in the HAT mutant. It is possible that the WT level of histone acetylation is sufficient for normal FACT occupancy, suggesting that histone acetylation, but not deacetylation, plays a role in FACT recruitment/retention in coding regions. As such, any further increase does not necessarily increase FACT occupancy. Taken together, these results suggest a role for the acetylated H3 tail in promoting FACT occupancy in the transcribed coding regions.

**Figure 2:**
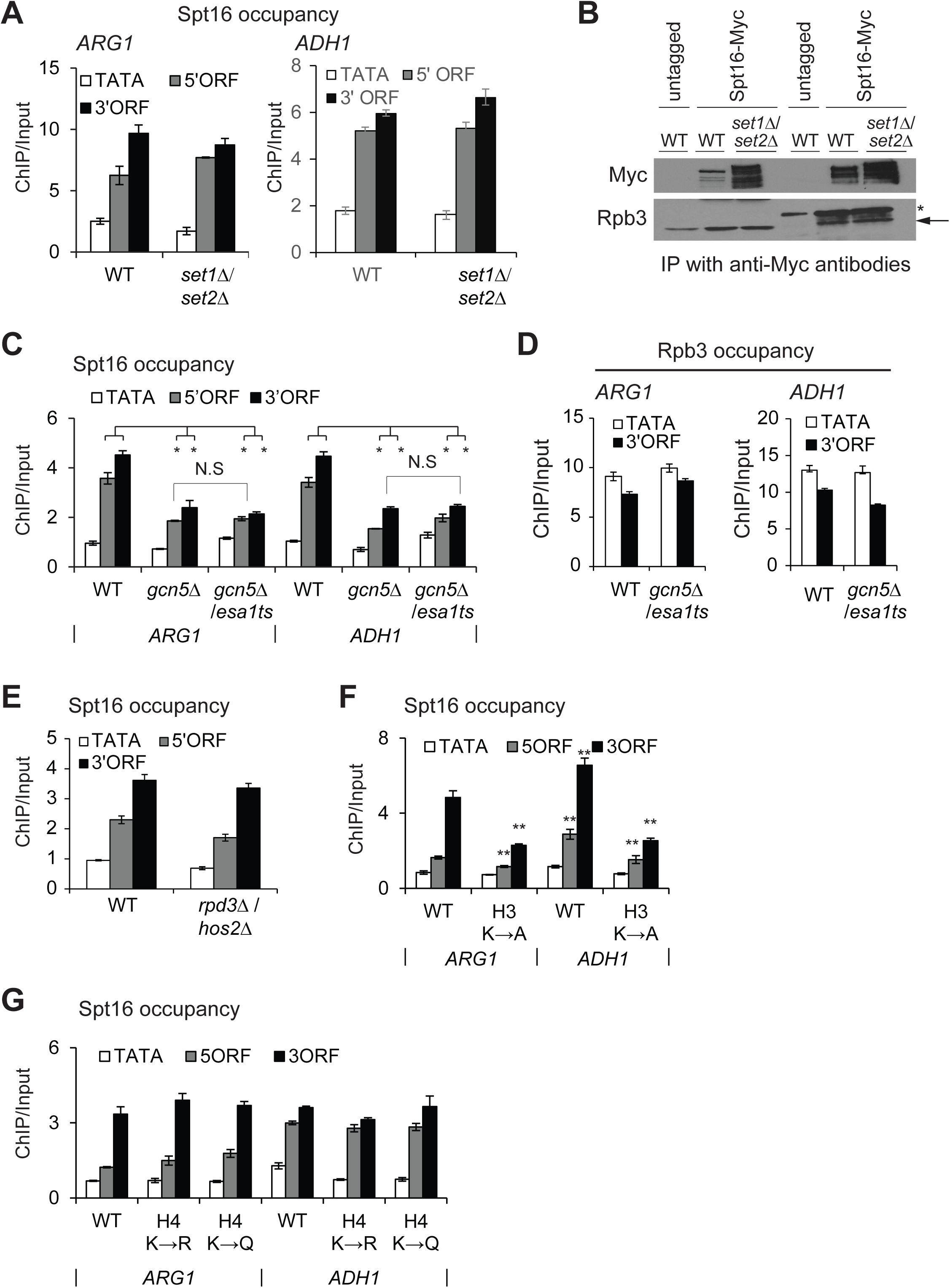
Role of the acetylated H3 tail in modulating Spt16 occupancy. A) ChIP enrichment of Spt16-Myc at *ARG1* in WT and histone methyltransferase mutant *set1*Δ/*set2*Δ. * represents a p-value < 0.001, and ** represents a p-vlaue <0.0001. B) Whole-cell extracts prepared from Spt16-Myc tagged WT and *set1Δ/set2Δ* strains were immunoprecipitated with anti-Myc antibodies, and the immunoprecipitates were analyzed by western blot to detect signals for Myc and Rpb3. Untagged WT was used as a control. Rpb3 band in immunoprecipitated samples is shown by an arrow and the IgG heavy chain by an asterisk (*). C) Spt16-Myc ChIP occupancy in WT and *gcn5*Δ/*esa1* histone acetyletransferase (HAT) mutant. Spt16-Myc occupancies were measured by ChIP at *ARG1* and *ADH1* in a *gcn5*Δ/*esa1ts* strain grown either at 25°C (represented as *gcn5*Δ) or at 37°C to inactivate Esa1 (*gcn5*Δ/*esa1*). Graphs show mean and SEM. * represents a p-value < 0.001, and ** represents a p-vlaue <0.0001. Spt16 occupancy differences between *gcn5*Δ and *gcn5*Δ/*esa1ts* were not significnat (N.S). D) Rpb3 occupancies in WT and *gcn5*Δ/*esa1ts* at *ARG1* and *ADH1*. E) Spt16 occupancies in WT and HDAC mutant *hos2*Δ/*rpd3*Δ at *ARG1*. F-G) Spt16-Myc enrichments at *ARG1* and at *ADH1* in histone H3 (F) and H4 tail mutants (G). H3 K4, K9, K14, K18 substituted to alanine, H3K→A; H4 K5, K8, K12 and K16 substituted to arginine; H4K→R, or to glutamine H4K→Q. Graphs show mean and SEM. ** represents a p-vlaue <0.0001.

To provide additional proof for the role of histone acetylation, we examined Spt16 occupancy in the H3 and H4 tail point mutants. The H3 mutant (K4, K9, K14, K18 substituted to alanine; H3K→A) displayed reduced Spt16 occupancy in the ORFs of both *ARG1* and *ADH1* genes (Figure 2F). However, only minimal changes in Spt16 occupancy were observed in the H4 mutant (K5, K8, K12 and K16 substituted to arginine; H4K→R, or to glutamine H4K→Q) (Figure 2G; H4K→A mutant exhibits a lethal phenotype). While Spt16 interacts with both H3 and H4 N-terminal tails, and the histone tail mutants impair FACT function (BISWAS *et al*. 2006; VANDEMARK *et al*. 2008), our results suggest that acetylation of H3 tail makes a greater contribution to the FACT occupancy in the coding regions.

### FACT and Spt6 are required for transcription genome-wide

Microarray analyses examining transcription defects have revealed that the loss of Spt16 function leads to aberrant transcription at many genomic locations, including within coding regions (CHEUNG *et al*. 2008; VAN BAKEL *et al*. 2013). While it is evident that Spt16 functions to suppress wide-spread cryptic and anti-sense transcription, the role of FACT in regulating Pol II occupancy in coding regions is not well understood at a genome-wide scale. Gene-specific studies have suggested that FACT regulates transcription at the initiation step (BISWAS *et al*. 2006; JIMENO-GONZALEZ *et al*. 2006; DUINA *et al*. 2007). Additionally, FACT, in cooperation with H2B ubiquitination, is important for restoration of chromatin in the wake of Pol II elongation (FLEMING *et al*. 2008). As mentioned earlier, similar to the FACT complex, Spt6 (a H3/H4 chaperone) is localized to the coding regions of strongly transcribed genes (MAYER *et al*. 2010; IVANOVSKA *et al*. 2011; PERALES *et al*. 2013; BURUGULA *et al*. 2014) and is also important for suppressing aberrant transcription (KAPLAN *et al*. 2003; CHEUNG *et al*. 2008; VAN BAKEL *et al*. 2013). To compare the impact of FACT and Spt6 on transcription, we utilized strains in which the expression of *SPT16* or *SPT6* was under the control of a tetracycline repressible promoter (*SPT16-TET* and *SPT6-TET*). These promoters can be repressed by growing cells in the presence of doxycycline (dox).

To rule out unexpected consequences of replacing the endogenous promoter with the TET-promoter, we first compared Spt16 and Spt6 protein levels in TET-strains and BY4741 (*S. cerevisiae* WT strain). Spt16 and Spt6 protein levels in the untreated (no dox; ND) *SPT16-TET* and *SPT6-TET*, respectively, were very similar to those detected in the BY4741 cells (Figure 3A, top panel). As expected, treating *SPT16-TET* and *SPT6-TET* cells with dox led to reduced expression of Spt16 and Spt6, respectively (Figure 3A, bottom panel). We noted that Spt16 was depleted to a greater extent than Spt6 upon dox-treatment. Since Spt16 mutants have been shown to cause cell cycle defects (PRENDERGAST *et al*. 1990), we also measured the level of budded and unbudded cells in BY4741, *SPT16-TET* and *SPT6-TET* dox-treated cells. We did not find any significant increase in the number of budded and unbudded cells under Spt6 or Spt16 depleted conditions (Figure S3A), suggesting that depleting Spt16 or Spt6, under the experimental conditions employed, elicit minimal cell cycle defects. Moreover, the TET-strains grown in the presence or absence of dox, exhibited similar viability (Figure 3B). Altogether, these results indicate that the untreated *SPT16-TET* and *SPT6-TET* cells behave similar to BY4741.

**Figure 3:**
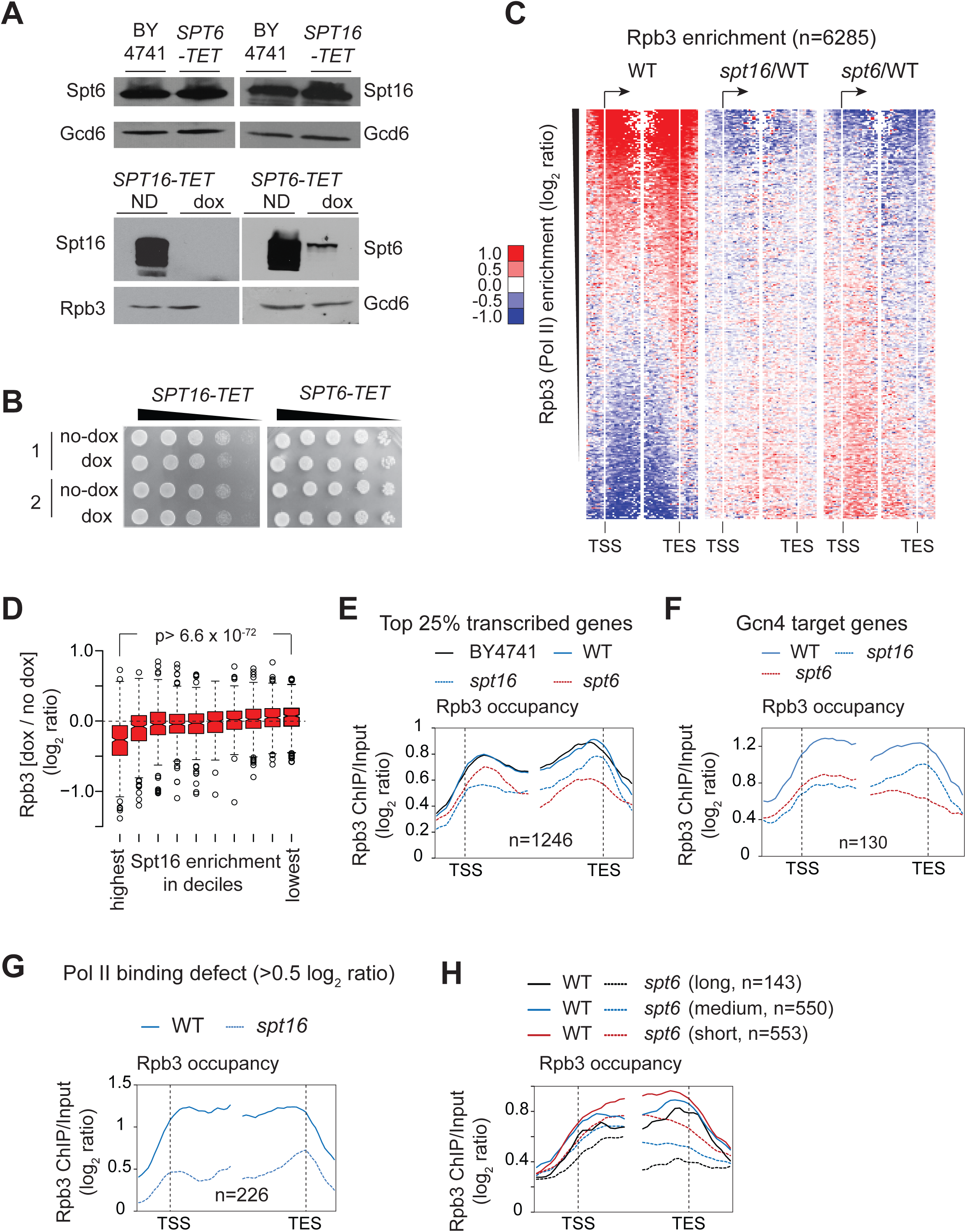
Effect of depleting Spt16 and Spt6 on Pol II occupancy genome-wide. A) *BY4741*, *SPT16-TET* and *SPT6-TET* cells were grown in SC media and induced by SM. Western blots show Spt16 and Spt6 protein levels in BY4741, *SPT16-TET* and *SPT6-TET* strains without dox-treatment (top panel). The Spt16 and Spt6 levels in untreated (ND) and dox-treated TET-strains are shown (bottom panel). B) *SPT16-TET* and *SPT6-TET* cells grown and subcultured in SC media, with and without doxycycline (dox), were collected, serially 10-fold diluted, and spotted on SC plates. Growth for the two cultures of each *SPT16-TET* (left) and *SPT6-TET* (right) with and without dox (no-dox) treatment are shown. C) Heat-maps depicting genome-wide Rpb3 (Pol II) enrichment in *SPT16-TET* (without dox; WT) (left), and changes in Rpb3 occupancies in dox-treated *SPT16-TET* (*spt16*/WT) (middle) and *SPT6-TET* (*spt6*/WT) (right). Genes were sorted from highest to lowest ORF Rpb3 enrichment in WT cells. D) Box-plot showing the changes in Rpb3 occupancy according to the Spt16 enrichment in deciles. The average ORF occupancy of Rpb3 in *SPT16-TET* dox-treated and untreated cells was determined, and the change in Rpb3 occupancy on depleting Spt16 (dox/no dox) was calculated for each gene. Genes were then grouped into deciles according to the Spt16 occupancy in WT cells. The first decile shows the highest Spt16 occupancy and the 10^th^ shows the lowest. E) Rpb3 occupancy profiles for the top 25% Pol II-occupied genes (n=1246) in untreated *BY4741* and *SPT16-TET* (WT), and in dox-treated *SPT16-TET* (*spt16*) and *SPT6-TET* (*spt6*) cells are shown. F) Rpb3 occupancy profiles in untreated *SPT16-TET* (WT), and dox-treated *SPT16-TET* (*spt16*) and *SPT6-TET* (*spt6*) cells at the Gcn4 targets genes enriched in the top 25% Pol II-occupied genes. G) Rpb3 occupancy profile at the genes eliciting reduction in Rpb3 occupancy ≥ 0.5 log_2_ ratio (ChIP/input) on depleting Spt16 in WT and *spt16*. H) The top 25% Pol II-occupied genes were grouped on the basis of their gene-length and Rpb3 occupancy profiles for the long (>2 kb), medium (1-2 kb), and short (0.5-1 kb) are shown in the WT and Spt6-depleted cells.

To examine the effects of depleting Spt16 and Spt6 on transcription, we determined Rpb3 occupancy in untreated cells (*SPT16-TET*; referred to as WT hereafter), and dox-treated *SPT16-TET* (*spt16*) and *SPT6-TET* (*spt6*) cells by ChIP-chip. We observed a strong correlation (Pearson correlation, r= 0.93) between the Pol II occupancies in the dox-untreated *SPT16-TET* and BY4741 strains, genome-wide, which further indicated that replacing the endogenous *SPT16* promoter with the TET-promoter does not adversely affect transcription. We also determined Spt16 occupancy genome-wide by ChIP-chip and found that Spt16 occupancy in coding regions strongly correlated with Pol II occupancy (Pearson correlation, r=0.85), in agreement with previous studies (MAYER *et al*. 2010).

The heat-maps depicting changes in Rpb3 enrichment (*spt16*/WT) showed diminished ratios in coding regions of the genes displaying the greatest Rpb3 enrichments in WT cells (Figure 3C). Consistent with a strong correlation between Spt16 and Rpb3 occupancies, the genes with the highest Spt16 enrichments showed greatest Rpb3 reductions (Figure 3D). We also noticed that depletion of Spt16 (and Spt6, described later) also revealed an increase in Pol II occupancies at those genes, which otherwise show very poor enrichment ratios in WT cells. Given that our ChIP-chip normalization was performed without spike-in control, it is difficult to ascertain the apparent increase in Rpb3 occupancy as biologically relevant.

To further analyze the impact of depleting Spt16 on Pol II occupancy, we selected the top 25% genes showing greatest Rpb3 occupancy in WT cells (n=1246). Nearly identical profiles for Rpb3 occupancy were observed in WT and BY4741 at the metagene comprised of these transcribed genes (Figure 3E). In contrast, Spt16 depletion evoked reduction in Pol II occupancy in the coding region of these sets of genes. Given that Gcn4 target genes are activated under the growth conditions used (see Materials and Methods), we additionally analyzed the effect of depleting Spt16 on transcription of Gcn4 targets. 130 Gcn4-regulated genes were enriched among the top 1246 transcribed genes. Reduced Pol II occupancies were evident in coding regions of these genes as well in Spt16-depleted cells (Figure 3F). Further, we identified 226 genes showing reduction in Pol II occupancy ≥0.5 log_2_ ratio (ChIP/input) in Spt16-depleted cells (Figure 3G). Interestingly, these genes were enriched among the top 10% transcribed genes (p-value=10^−117^), suggesting that depletion of Spt16 imparts a significant effect on Pol II occupancy at highly expressed genes.

We noticed a small increase in Pol II occupancy in the Spt16-depleted cells at the 3’ ends of genes, analyzed above (Figure 3E). This increase may reflect increased cryptic transcription events that accumulate Pol II from the 5’ to 3’ end. Such an explanation would be consistent with the established role of histone chaperones in suppressing cryptic transcription (KAPLAN *et al*. 2003; CHEUNG *et al*. 2008; VAN BAKEL *et al*. 2013). A previous study utilizing microarray predicted 960 and 1130 genes to have cryptic transcription in *spt6-1004* and *spt16-197* mutants (CHEUNG *et al*. 2008). We found that only 154 genes, predicted to express cryptic transcripts in the previous study, were among the 1246 genes exhibiting high-levels of Pol II occupancy (Figure S3B). Replotting Pol II occupancy data after excluding these 154 genes (Figure S3C) displayed profiles similar to that observed in Figure 3E. Pol II pausing and queuing in the 3’ end superimposed on elongation defects at the very 5’ end could potentially explain an apparent increase in Pol II occupancy towards the 3’ end on depleting Spt16. Regardless, our results showing reduction in Pol II occupancy across the coding regions is consistent with previous studies indicating a role for FACT in transcription initiation or early stages of Pol II elongation.

Next, we analyzed Pol II occupancy in the Spt6 depleted cells. Interestingly, at the top 25% of Pol II-occupied genes, Spt6-depletion elicited a greater reduction in Rpb3 occupancy towards the 3’ end (Figures 3C (right) and 3E), consistent with previous studies showing the greatest Spt6 enrichments in the 3’ ends of transcribed genes (PERALES *et al*. 2013; BURUGULA *et al*. 2014). A 5’ to 3’ bias in Pol II occupancy was also evident at the Gcn4-targets (Figure 3F) and at 238 genes, which showed a reduction in Pol II occupancy ≥ 0.5 log_2_ ratio (ChIP/input) (Figure S3D). A progressive reduction in Pol II occupancy in the 5’ to 3’ direction, in Spt6-depleted cells, suggests that Spt6 may regulate Pol II processivity. Alternately, given the role of Spt6 activity in the 3’—mRNA processing, diminished Spt6 levels could also result in reduced Pol II occupancy at the 3’ end (KAPLAN *et al*. 2005). To distinguish between these two possibilities, we analyzed Rpb3 occupancy at the top 25% transcribing genes based on their gene length. All three groups of genes, long (> 2 kb), medium (1-2 kb) and short (0.5-1 kb), showed a 5’ to 3’ bias in Pol II occupancies (Figure 3H). Interestingly, however, long and medium genes displayed a greater reduction in the 3’ end compared to the short genes (0.5-1 kb) (Figure 3H and S3E). A simpler explanation for this observation is that Spt6 promotes Pol II processivity, in agreement with previous studies (ENDOH *et al*. 2004; ARDEHALI *et al*. 2009; PERALES *et al*. 2013). A 3’—mRNA processing defect (KAPLAN *et al*. 2005) would be expected to produce similar reductions in Pol II occupancies at the 3’ end irrespective of gene length. Collectively, our data suggest that loss of FACT and Spt6 functions produces distinct effects on Pol II occupancy. However, the role of FACT and Spt6 in suppressing aberrant transcription, to some extent, could have an effect on Pol II occupancy under depletion conditions.

### FACT and Spt6 differentially impact transcription and histone occupancy

To further address the functional overlap between FACT and Spt6, we examined genes, which showed a reduction in Pol II occupancy (≤ −0.5 log_2_ ratio) upon depleting these factors. We found a significant overlap between the genes exhibiting Pol II fold-change log_2_ ≥ 0.5 upon depleting either Spt16 or Spt6 (p-value = 5.1 × 10^−76^, n= 111) (Figure 4A). The genes showing Pol II occupancy defects upon depleting either Spt16 or Spt6 (common; n=111) exhibited, on average, higher Pol II occupancy (in WT cells) than those genes which showed defects only after depleting either Spt16 or Spt6 (unique) (Figure 4B). This observation suggests that strongly transcribed genes may need full functions of Spt6 and Spt16 for a high-level of transcription. It is also interesting to note that while Spt6 depletion was less efficient compared to that of Spt16, it nonetheless evoked very similar Pol II occupancy defects on these genes (Figure 4C). Rpb3 profiles at the genes uniquely affected either by Spt6 or Spt16 depletion showed expected profiles (Figures 4D and 4E). Considering that Spt6 exhibits higher occupancy towards the 3’ ends (MAYER *et al*. 2010; PERALES *et al*. 2013; BURUGULA *et al*. 2014), we analysed distribution of the gene-length in the three classes of genes. The longer genes were enriched in the Spt6-unique and common genes, whereas Spt16-unique genes were shorter in comparison (Figure S4A and S4B). Thus, it appears that the differential effect of Spt16 and Spt6 depletion could partly be due to differences in their localization patterns over the coding sequences and transcription level in WT conditions.

**Figure 4:**
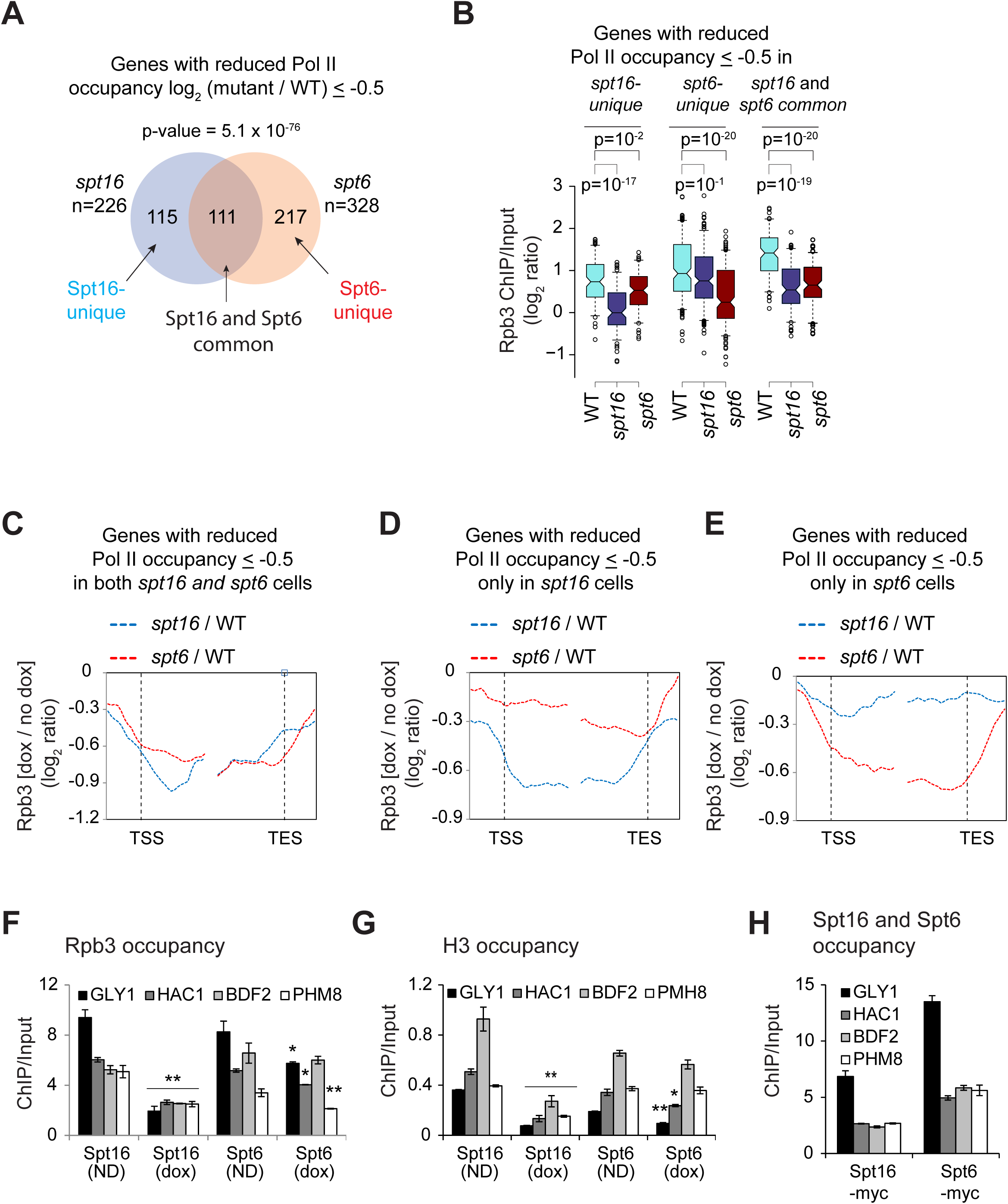
FACT promotes nucleosome reassembly. A) Venn diagram showing overlap among the genes which displayed Rpb3 occupancy defect >0.5 log_2_ ratio (ChIP/Input) in Spt16 and Spt6 depleted cells. p-value for the overlap is shown. B) Box-plot showing Rpb3 average ChIP-chip enrichments at Spt16-unique, Spt6-unique, and Spt16 and Spt6 common genes in WT, Spt16-depleted and Spt6-depleted cells. C-E) Metagene analysis for changes in Pol II enrichment under Spt16 and Spt6 depeleted cells, for genes showing reduction > 0.5 log_2_ ratio (ChIP/Input) under Spt16 or Spt6 depleted condition (C; common), only under Spt16 depletion (D) only under Spt6 depletion (E). F-H) ChIP occupancies of Rpb3 (F), histone H3 (G), and of Spt6 and Spt16 in 5’ ORFs of indicated genes in *SPT16-TET* and *SPT6-TET* untreated (ND) or dox-treated (dox) cells. * represents a p-value < 0.05, and ** represents a p-vlaue <0.001.

We further examined the differential effect of depletion by determining Rpb3 ChIP occupancy at four genes, which showed comparable Spt6, and Spt16 occupancy in our ChIP-chip experiments. Pol II occupancies in the ORFs of *GLY1*, *HAC1*, *BDF2*, and *PHM8* were substantially reduced (∼2-5 folds) upon Spt16 depletion (Figure 4F). By contrast, only a moderate to negligible reduction was observed after depleting Spt6. For example, Rpb3 was reduced by less than 1.5 folds at *GLY1*, *HAC1*, *PMH8* upon Spt6 depletion. Similarly, we found that histone H3 occupancy was more severely reduced in 5’ ORFs of these genes upon depleting Spt16 than upon Spt6 (Figure 4G). Reduced H3 occupancies upon depletion of Spt16 and Spt6 are consistent with their role in histone reassembly (IVANOVSKA *et al*. 2011; PERALES *et al*. 2013; JERONIMO *et al*. 2015). Bigger reductions in Pol II and H3 occupancies were not due to higher Spt16 occupancies at these genes compared to that of Spt6 (Figure 4H). These analyses illustrate the importance for both FACT and Spt6 in regulating transcription globally. The differences observed on a subset of genes under the conditions of diminished FACT or Spt6 function might be a reflection of their apparent role in regulating different aspects of transcription, as well as on their different localization patterns across the gene.

### FACT cooperates with Spt6 to promote transcription

To further investigate whether Spt6 and FACT cooperate in promoting transcription, we deleted the Spt6 tandem SH2 domain (tSH2; 202 residues from C-terminus), which mediates Spt6 recruitment, genome-wide (MAYER *et al*. 2010; BURUGULA *et al*. 2014), in the *SPT16-TET* background (*SPT16/spt6*Δ*202*). As expected, treating the *SPT16/spt6*Δ*202* mutant with dox resulted in reduced Spt16 protein levels (Figure S5A). We then determined Rpb3 occupancies by ChIP-chip in untreated (S*PT16/spt6*Δ*202*) and dox-treated (*spt16/spt6*Δ*202*) cells, and compared the occupancies with that of the untreated *SPT16-TET* cells (WT). The changes in Rpb3 occupancy in untreated *SPT16/spt6*Δ*202* were significantly anti-correlated with the Rpb3 occupancy in the WT cell (r= −0.81) (Figure 5A), indicating a strong requirement of Spt6 in transcription, genome-wide. The changes in Pol II occupancies in the dox-treated *SPT16/spt6*Δ*202* showed very similar reductions as observed for the untreated cells. However, a modest but statistically significant reduction in Rpb3 occupancy was observed in dox-treated (*spt16/spt6*Δ*202)* cells compared to the untreated cells only for the first decile (p =5.2 × 10^−59^) representing the most highly transcribed genes (Figure 5B). This is consistent with the idea that both Spt6 and Spt16 are necessary to elicit high transcription levels.

**Figure 5:**
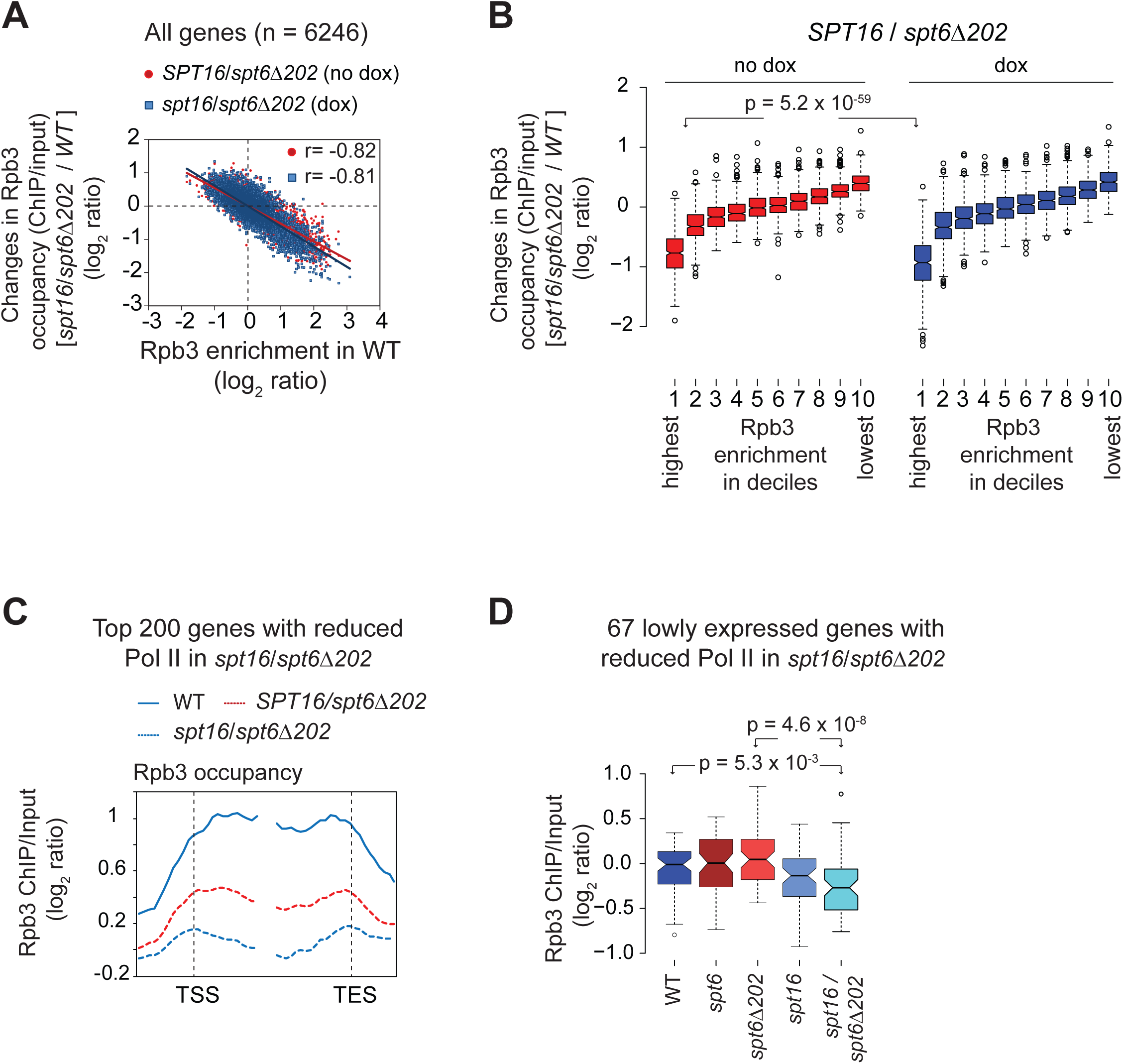
Spt6 promotes histone eviction and transcription. A) The changes in Rpb3 occupancies (log_2_ ratio, ChIP/input) in the untreated (no dox) and treated (dox) *spt16/spt6*Δ*202* cells relative to occupancies in the untreated *SPT16-TET* (WT) are plotted against the Rpb3 ORF occupancies in WT. Pearson correlations are shown. B) Box-plot showing changes in Rpb3 occupancy in *SPT16/spt6*Δ*202* (dox and no dox) relative to the *SPT16-TET* (no dox) WT cells, at the genes grouped in deciles on the basis of the average ORF Rpb3 occupancies observed in WT cells. The first decile shows the highest Rpb3 occupancy and the 10^th^ shows the lowest. C) Top 200 genes showing the greatest reduction in Pol II occupancy in dox-treated *SPT16/spt6*Δ*202* relative to the untreated cells were selected based on the averge Rpb3 enrichments in coding regions. Pol II occupancy profile for these genes is plotted for WT, *SPT16/spt6*Δ*202* and *spt16/spt6*Δ*202*. D) Box-plot showing average Rpb3 enrichment (log_2_ ratio, ChIP/input) for 67 genes of the 200 genes. These genes were not among the top 25% genes showing greatest Rpb3 enrichment in WT cells. p-values are shown.

Rpb3 profiles in *spt16/spt6*Δ*202* mutant revealed greatly diminished occupancy across the coding regions of the top 25% Pol II-occupied genes (Figure S5B). It appeared that untreated *SPT16/spt6*Δ*202* elicited a greater reduction in Pol II occupancy than observed upon depleting Spt6 (compare Figures S5B and 3E). An incomplete depletion of Spt6 (Figure 3A), or different experimental conditions (no dox in *spt6*Δ*202* vs dox-treatment for Spt6 depletion) could account for this observation. Nonetheless, these results indicate the importance of Spt6 in stimulating high-level transcription, genome-wide. However, depleting Spt16 in *SPT16/spt6*Δ*202* cells produced only modest reductions in Pol II occupancy at the top 25% Pol II occupied (Figure S5B). This observation raises a possibility that FACT may require certain aspects of Spt6 function to stimulate transcription. Such an explanation is consistent with the observation that Spt16 depletion elicited reduced Pol II occupancy in otherwise WT strain (Figures 3D-G).

To further address functional cooperation between Spt16 and Spt6, we focused on the genes eliciting greater reduction in dox-treated *spt16/spt6*Δ*202* than in untreated cells. The top 200 genes showing the greatest Pol II occupancy defect revealed that depleting Spt16 in *spt6*Δ*202* background significantly reduced Rpb3 occupancy across the coding region (Figure 5C). 137 of these 200 genes were among the top 25% expressed genes (n=1246) in WT cells. The double mutant exhibited greater reduction in Pol II occupancy than observed in *spt6*Δ*202* or *spt16* cells at these 137 genes (Figure S5C). These observations support the idea that both FACT and Spt6 are needed for eliciting high-level of transcription. At the 63 lowly expressed genes among the 200 genes, significant reduction in Pol II occupancy was seen only in the double mutant *spt16/spt6*Δ*202* (Figure 5D). This could suggest that FACT acts redundantly with Spt6 in promoting transcription of a subset of lowly expressed genes. More sensitive methods will be required to fully comprehend the extent to which Spt6 and FACT coordinate transcription of lowly expressed genes, especially considering the technical challenges in accurately measuring Pol II occupancies at genes expressed at very low levels.

## DISCUSSION

In this study, we have examined the role of two highly conserved histone chaperones, Spt16 and Spt6, in regulating genome-wide transcription, under amino acid starvation conditions. Spt6 recruitment to coding regions is stimulated by the phosphorylated Pol II CTD and by HDACs Rpd3 and Hos2 (MAYER *et al*. 2012; BURUGULA *et al*. 2014). However, the mechanism by which the FACT complex associates with chromatin is not well understood. Our results showing diminished occupancy of Spt16 at the ORFs of several genes and reduced interaction with nucleosomes in the H3 tail mutant (Figure 1), suggests a role for the H3 tail in promoting FACT association with transcribed regions. Our results additionally suggest that histone acetylation enhances the ability of the FACT complex to associate with transcribed ORFs, since deleting Gcn5 (a H3 HAT) or mutating the H3 tail lysine residues significantly dampened Spt16 enrichment in coding sequences of *ARG1* and *ADH1* (Figures 2C and 2F). These results provide *in vivo* evidence for the previous studies showing impaired FACT binding to the nucleosomes/histones lacking N-terminal tails (VANDEMARK *et al*. 2008; WINKLER *et al*. 2011). Currently, however, which regions of Spt16 are required for histone tail interactions is unclear. While Spt16 N-terminal domain of *S. pombe* was shown to be important for interacting with H3 and H4 tails (STUWE *et al*. 2008), in *S. cerevisiae*, this domain was found to be dispensable (VANDEMARK *et al*. 2008).

Considering that Spt16 exhibits similar affinity towards the acetylated and unacetylated histone peptides *in vitro* (STUWE *et al*. 2008), it is plausible that acetylation-mediated changes to the nucleosome structure allow FACT to stably bind chromatin at transcribing loci. Consistent with this idea, Gcn5 promotes histone eviction, Pol II elongation, and stimulates recruitment of bromodomain-containing chromatin remodelers, RSC and SWI/SNF (GOVIND *et al*. 2007; DUTTA *et al*. 2014; SPAIN *et al*. 2014). Additional contacts with core domains of H3/H4 and with H2A/H2B through its C-terminal domain (WINKLER *et al*. 2011; KEMBLE *et al*. 2015) could further stabilize FACT-chromatin interactions, thereby maintaining chromatin in an accessible conformation to promote transcription, and concomitantly aiding in reassembly of evicted histones in the wake of transcription (JAMAI *et al*. 2009). Additional factors may act cooperatively with other factors to enhance FACT enrichment. For instance, FACT interacts with the Paf1 complex, and chromatin remodeler Chd1, both of which are enriched in transcribed regions (KROGAN *et al*. 2002; SQUAZZO *et al*. 2002; SIMIC *et al*. 2003).

Enrichment of FACT in coding regions (MAYER *et al*. 2010) (*data not shown*), and its ability to promote Pol II transcription through the nucleosomal templates, *in vitro*, (BELOTSERKOVSKAYA *et al*. 2003; HSIEH *et al*. 2013), strongly suggests a role for FACT in the elongation step of transcription. FACT is linked to the reestablishment of the disrupted chromatin structure in the wake of transcription (JAMAI *et al*. 2009), and consequently, in suppressing aberrant transcription by preventing utilization of cryptic promoters (KAPLAN *et al*. 2003; CHEUNG *et al*. 2008; VAN BAKEL *et al*. 2013). Our data indicate that FACT is globally required for promoting transcription in the coding regions. Spt16 deficiency reduced Pol II occupancy in coding regions of highly transcribed genes, including of the ribosomal protein genes and Gcn4 targets (Figure 3). It was previously reported that histone mutants which perturb association of Spt16 (shift in occupancy towards the 3’ end) also reduced Pol II occupancy at the 5’ end of *PMA1* and *FBA1* genes (NGUYEN *et al*. 2013). Thus, it seems that impairing FACT activity, through depletion of Spt16 or by altering FACT association in the coding region (NGUYEN *et al*. 2013), elicit greater defects in transcription in the early transcribed regions. These results raise a possibility that FACT may help polymerases in negotiating the nucleosomal barrier downstream of the TSS. Such an idea is consistent with biochemical studies showing that FACT relieves Pol II pauses well within the nucleosomes and acts primarily to overcome tetramer-DNA contacts (BONDARENKO *et al*. 2006). Considering that the rate of elongation in the early transcribed regions is slower than that observed in the mid or 3’ end regions (DANKO *et al*. 2013), our findings suggest that FACT might help polymerases to overcome the nucleosomal barrier at the 5’ ends of the transcribed genes. Although, nucleosomes are expected to pose a similar block to elongating Pol II irrespective of their position along coding sequences, nucleosomal impediment to transcription at the distal end may be additionally relieved by factors, such as chromatin remodelers RSC and SWI/SNF, which are also enriched in the ORFs of many genes (DUTTA *et al*. 2014; SPAIN *et al*. 2014). Moreover, post-translational histone modifications, which are not uniform across coding regions, may have a differential effect on the FACT activity to stimulate transcription. In support of such a possibility, it was shown that the FACT complex cooperates with H2B ubiquitination to regulate transcription both *in vitro* and *in vivo* (PAVRI *et al*. 2006; FLEMING *et al*. 2008).

In contrast to FACT, Spt6-depletion diminished Pol II occupancy at the 3’ end (Figures 3E-F and 3H). Lower Pol II occupancy towards the 3’ end could result from a processivity defect and/or from a specific defect at the 3’ end. Our analyses, based on gene-length, favor the idea that Spt6 enhances Pol II processivity, possibly through stimulating nucleosome eviction, or by preventing premature dissociation during Pol II traversal through coding regions. Such an interpretation is consistent with previous studies showing reduced Pol II occupancy at the 3’ end of exceptionally long-genes (PERALES *et al*. 2013). Spt6 has been also shown to enhance the rate of Pol II elongation at heat-shock genes in *Drosophila* (ARDEHALI *et al*. 2009). Therefore, reduced Pol II occupancy near the 3’ ends of yeast genes suggests that Spt6 may function to help elongating polymerase traverse coding regions through multiple mechanisms.

Pol II occupancy in the Spt6 mutant lacking the C-terminal tandem SH2 domain was strongly correlated with Pol II occupancy in the WT cells, indicating a strong requirement for Spt6 in stimulating transcription genome-wide. Even though depleting Spt16 in otherwise WT cells evoked reduction in Pol II occupancy from highly transcribed genes, this surprisingly had only a minor impact on Pol II occupancy at the majority of transcribed genes in *spt6*Δ*202* cells. This suggests that both factors might have overlapping roles and that FACT may rely on the certain aspects of Spt6 functions in promoting transcription. It makes sense considering that FACT, and Spt6 are recruited to transcribed regions (MAYER *et al*. 2010; BURUGULA *et al*. 2014), and that both factors can interact with H2A-H2B and H3-H4 (MCCULLOUGH *et al*. 2015). Despite interacting with nucleosomes as well as with histones, unlike Spt6, FACT can reorganize the nucleosome structure in a manner that increases DNA accessibility. This distinct ability of FACT could explain reduction in Pol II occupancy observed in a subset of transcribed genes in *spt16/spt6*Δ*202* double mutant, but not in the single mutant (Figure 5C). Overall, our study strongly implicates both chaperones in promoting transcription genome-wide.

## ACKNOWLEDGEMENTS

We thank Tim Formosa for providing antibodies against Spt6 and Spt16. We also thank Alan Hinnebusch, Randy Morse and Jeena Kinney for useful discussions and providing valuable comments on the manuscript. CKG is supported by grants from the National Institutes of Health (GM095514), and Center for Biomedical Research (CBR, Oakland University). We also acknowledge the genomic core facility at Michigan State University.

## Legends to the supplemental figures

**Figure S1:**
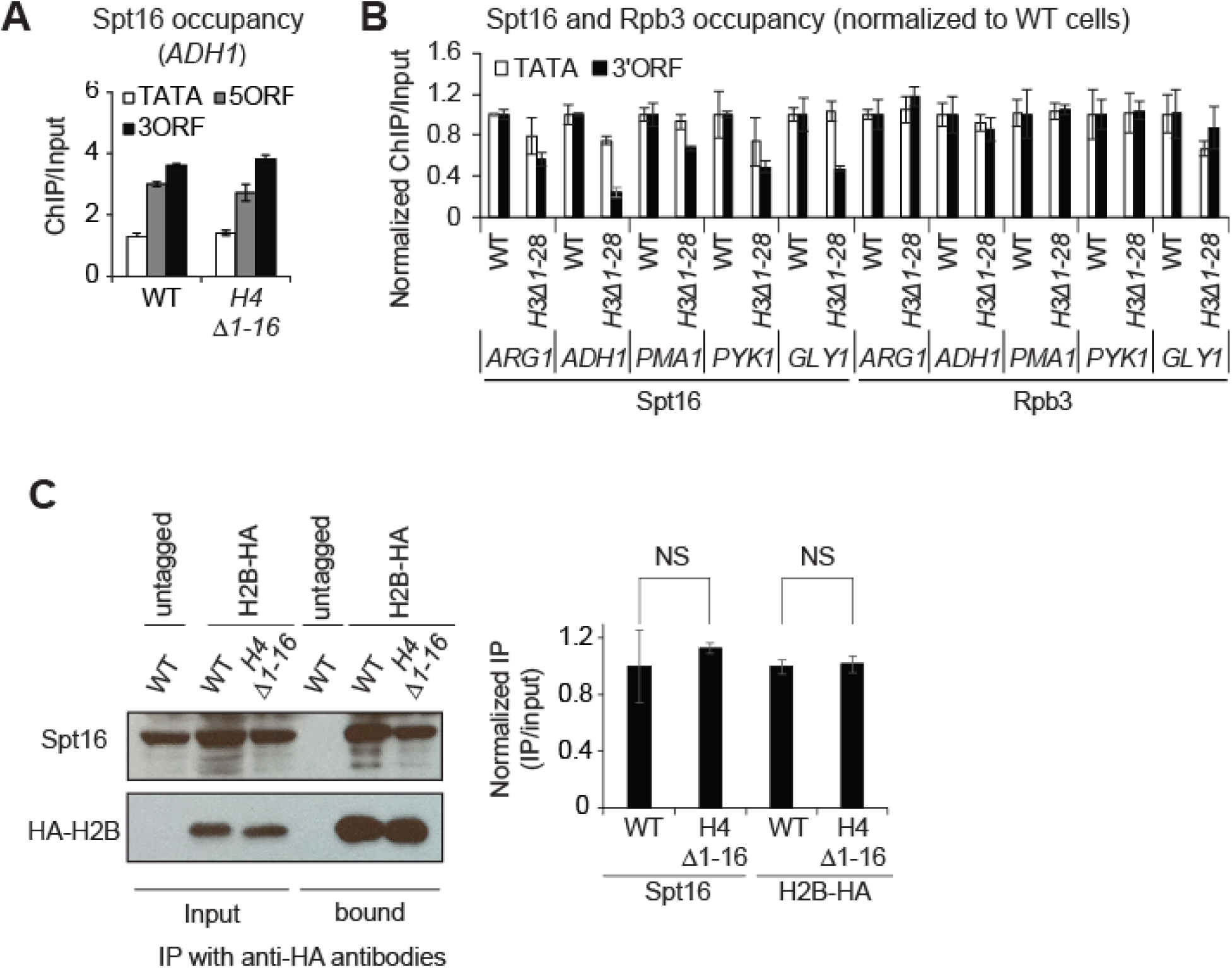
Role of the H3 tail in FACT recruitment. A) Spt16 ChIP occupancy at *ADH1* in a H4 tail mutant lacking the first 16 residues (*H4*Δ*1-16*). B) Spt16 and Rpb3 ChIP occupancy in the mutant was normalized against the WT occupancy at each gene and the normalized data is presented for the indicated genes. C) Anti-HA beads were used to pull down HA-tagged H2B from the whole cell extracts prepared from the untagged WT, the HA-tagged WT and *H4*Δ*1-16* cells. The immunoprecipitates were subjected to western blot using anti-HA and anti-Spt16 antibodies. The representative blot is shown on the left, and the quantified data on the right.

**Figure S2:**
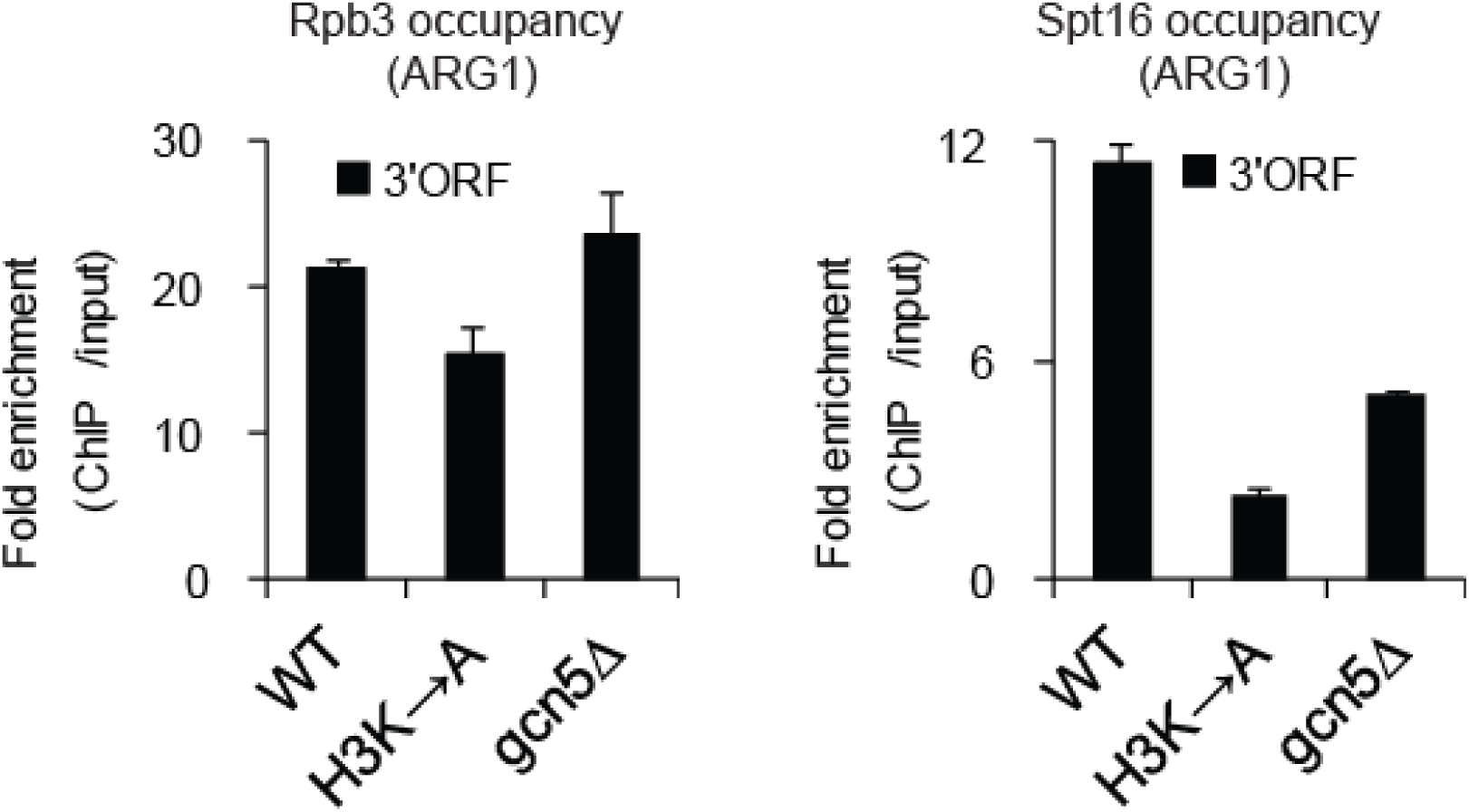
Role of histone acetylation in recruiting Spt16. ChIP occupancies of Rpb3 (left) and Spt16 (right) at *ARG1* in WT, *H3K*→*A*, and *gcn5*Δ mutants were determined by real-time (RT) PCR. ChIP occupancy in the *ARG1* 3’ ORF in WT were normalized to *POL1* signals from the same ChIP sample, and normalized again with the *ARG1/POL1* signals for the related input samples. Average data with SEM for three independent experiments is shown.

**Figure S3:**
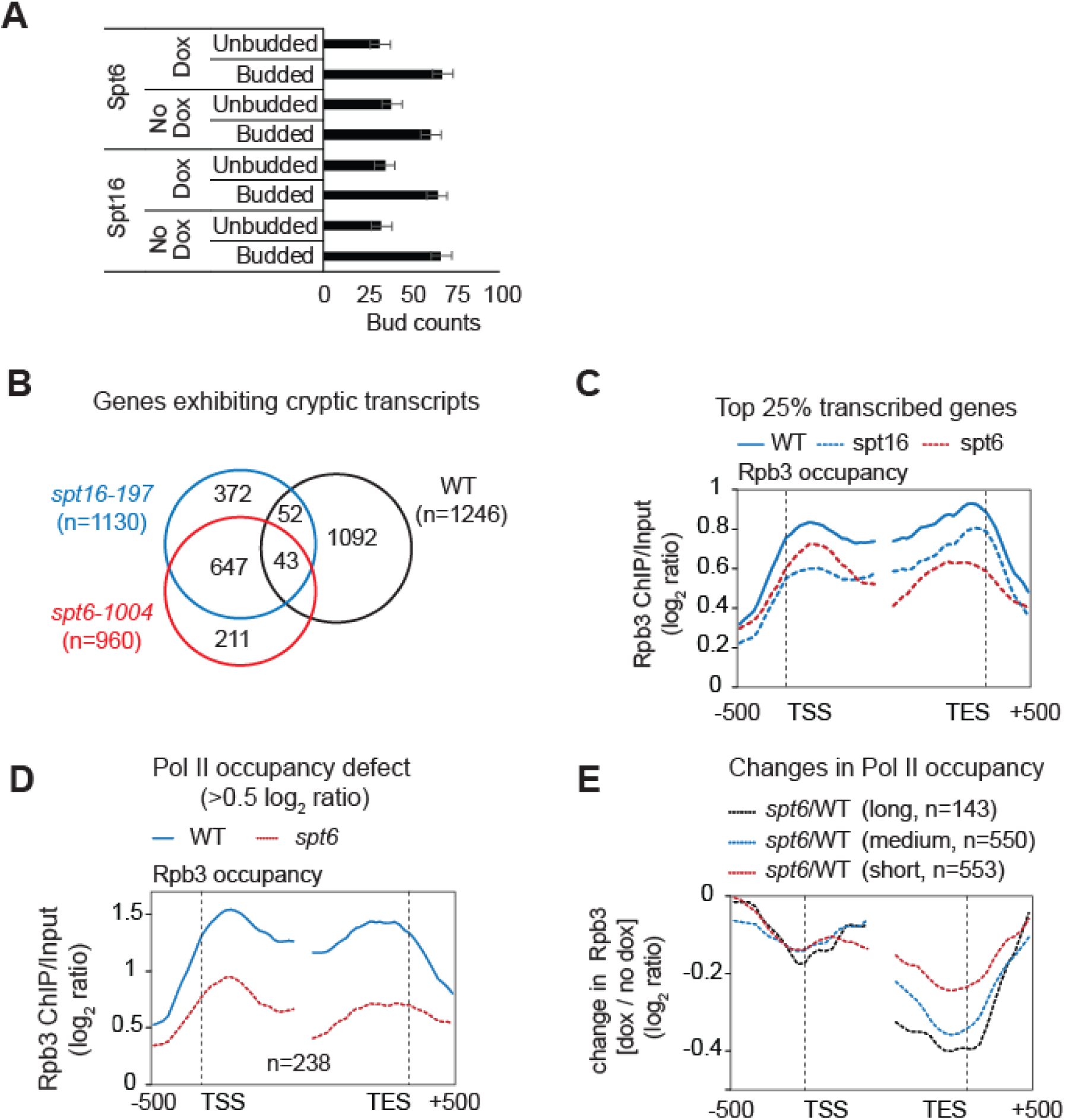
Rpb3 occupancy in WT, and Spt16 and Spt6 depleted cells. A) A bar-graph showing the bud counts in *SPT16-TET* and *SPT6-TET* cells in the presence or in the absence of doxycycline (dox). The cells were grown exactly as they were for the ChIP-chip experiments. The cells were briefly sonicated before counting budded and unbudded cells under the microscope. The error bars represent standard error of the mean. B) Venn diagram showing overlap between the top 25% Pol II-occupied genes and those genes which were identified to express cryptic transcripts in *spt16-197* and *spt6-1004* (Cheung et al. 2008). C) Overlapping genes, identified to express cryptic transcripts and were among the top 25% Pol II occupied genes (n=154), were removed from the list of 1246 highly expressed genes. Pol II occupancy profile for the remaining 1092 genes in WT, spt16 and spt6 cells is shown. D) Rpb3 occupancy profile of WT and *spt6* cells at the genes eliciting reduction in Rpb3 occupancy ≥ 0.5 log_2_ ratio (ChIP/input) upon depleting Spt6. E) The top 25% Pol II-occupied genes were grouped on the basis of their gene-length and changes in Rpb3 occupancy profiles (*spt6*/WT) for the long (>2 kb), medium (1-2 kb), and short (0.5-1 kb) are shown in the WT and Spt6-depleted cells.

**Figure S4:**
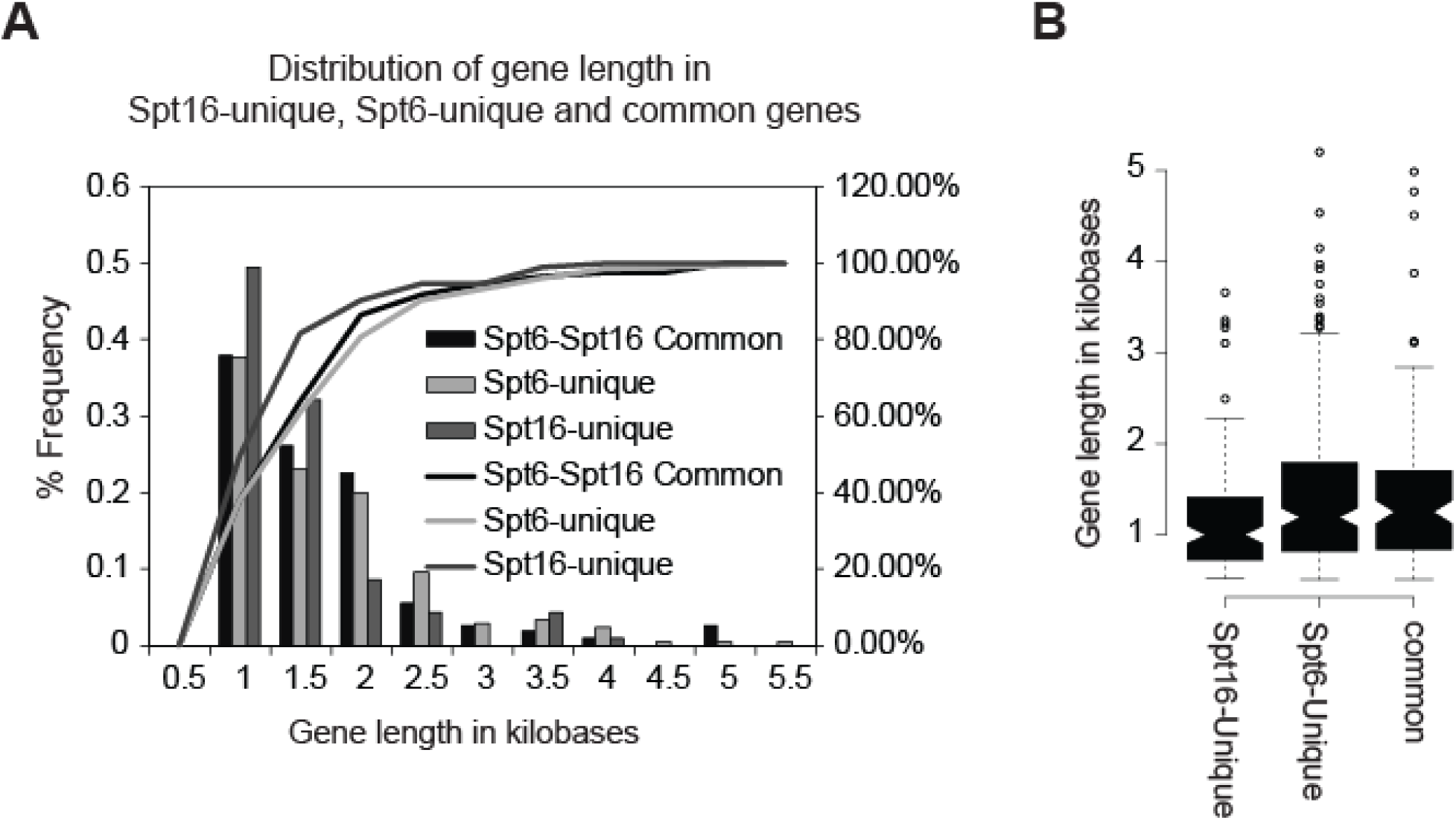
Gene-length analyses for the genes most affected by depletion of Spt6 or Spt16. A) The genes which showed more than 0.5 log_2_ reduction in Pol II occupancy upon depleting either Spt6 or Spt16 were analyzed, and were classified as Spt16-unique, Spt6-unique and Spt16-Spt6-common depending on whether they were enriched only upon depleting Spt16, Spt6 or both, respectively. Frequency distribution of these classes of genes based on their gene-length is plotted as a histogram. B) Gene-length for the three classes of genes is shown in the box plot. Center lines show the medians; box limits indicate the 25th and 75th percentiles as determined by R software; whiskers extend 1.5 times the interquartile range from the 25th and 75th percentiles, outliers are represented by dots. 115, 217 and 111 genes were present in the Spt16-unique, Spt6-unique and common gene-sets, respectively.

**Figure S5:**
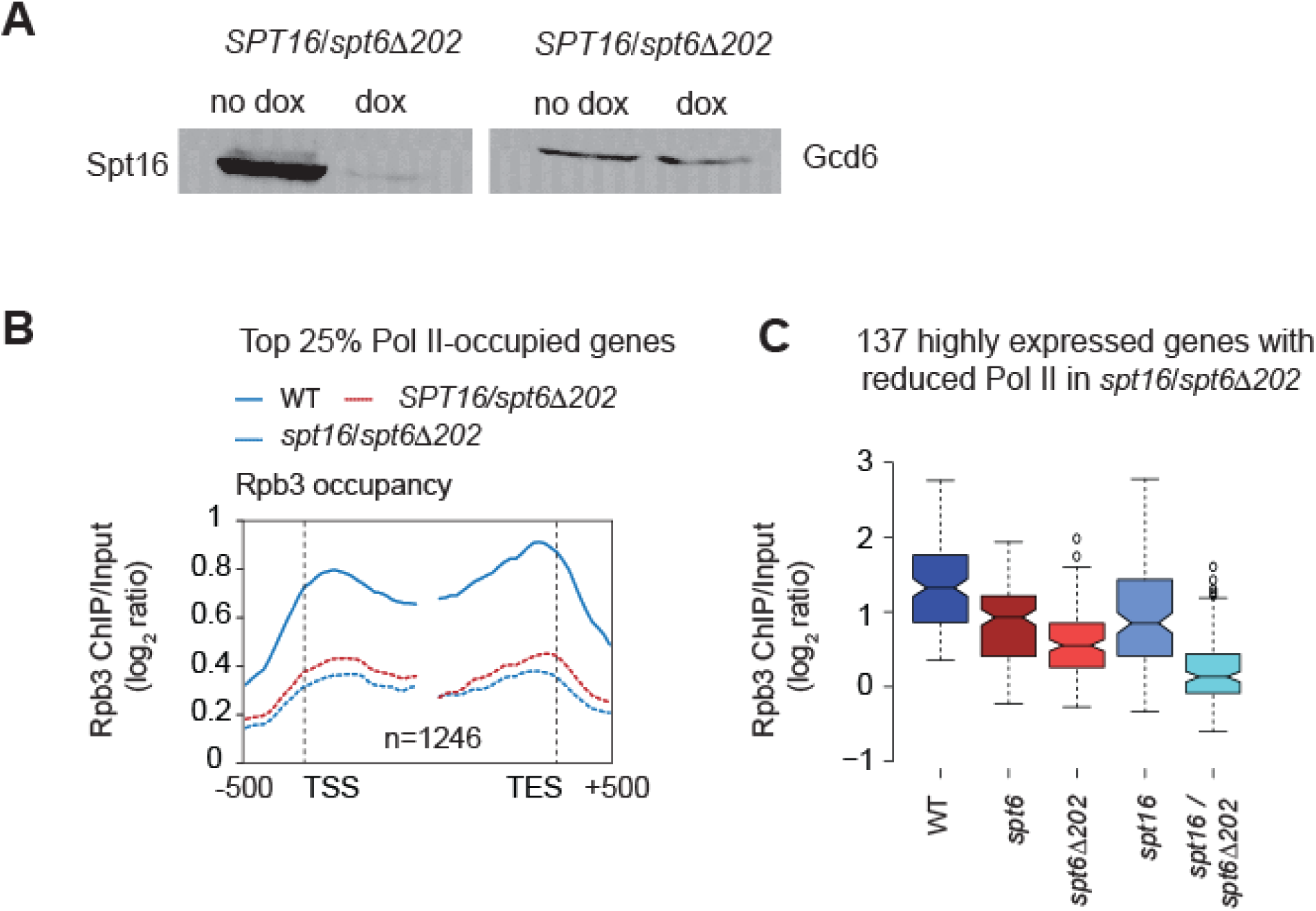
Pol II occupancy in *spt16/spt6*Δ*202* cells. A) Western blot showing Spt16 protein levels in untreated and dox-treated *Spt16/spt6*Δ*202* cells. B) Rpb3 occupancy profiles for the top 25% Pol II-occupied genes (n=1246) in untreated *SPT16/spt6Δ202* and dox-treated (*spt16/spt6*Δ*202*) cells. C) Box-plot showing Rpb3 enrichment (log_2_ ratio, ChIP/input) for the 137 highly expressed genes among the 200 genes showing greatest reduction in *Spt16/spt6*Δ*202* dox-treated compared to untreated cells. Pol II occupancies are shown for the untreated WT and *SPT16/spt6*Δ*202*, and dox-treated *SPT16-TET* (*spt16*) and *SPT6-TET* (*spt6*) and *SPT16/spt6*Δ*202* (*spt16/spt6*Δ*202*) cells.

**Table S1:**
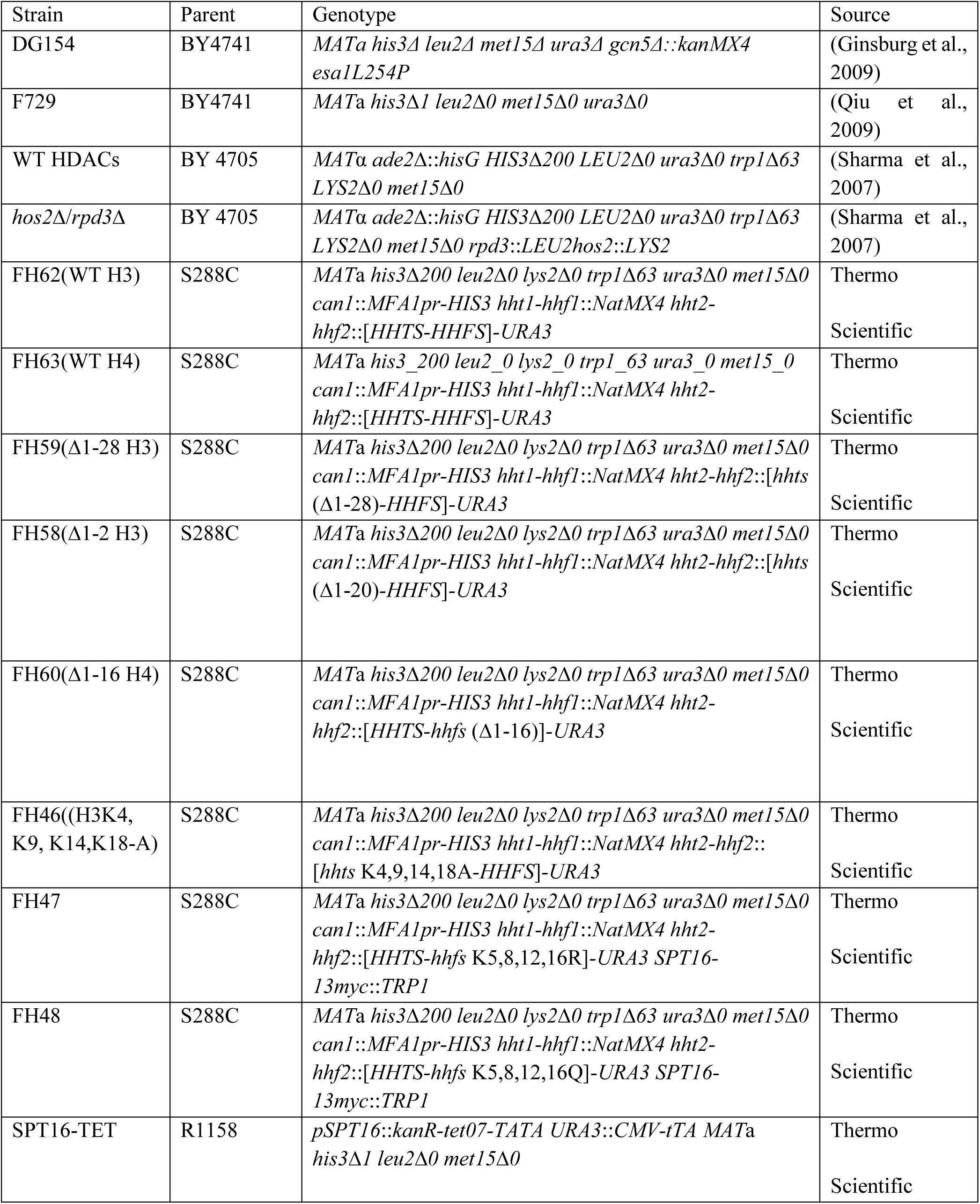
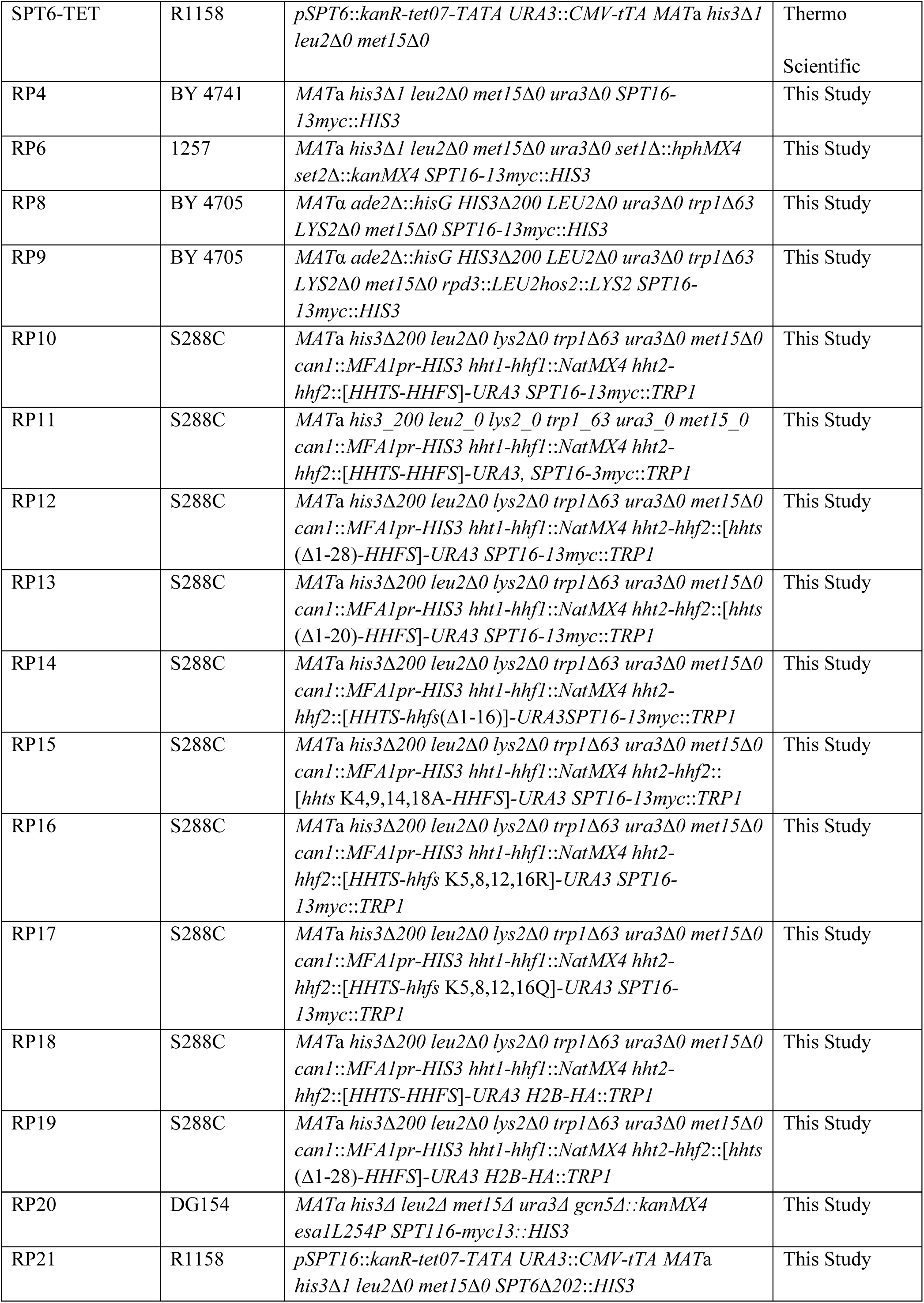
Yeast strains used in this study.

## REFERENCES

1. Andrulis, E. D., E. Guzman, P. Doring, J. Werner and J. T. Lis, 2000 High-resolution localization of Drosophila Spt5 and Spt6 at heat shock genes in vivo: roles in promoter proximal pausing and transcription elongation. Genes Dev 14: 2635–2649.

2. Ardehali, M. B., J. Yao, K. Adelman, N. J. Fuda, S. J. Petesch et al., 2009 Spt6 enhances the elongation rate of RNA polymerase II in vivo. EMBO J 28: 1067–1077.

3. Belotserkovskaya, R., S. Oh, V. A. Bondarenko, G. Orphanides, V. M. Studitsky et al., 2003 FACT facilitates transcription-dependent nucleosome alteration. Science 301: 1090–1093.

4. Billon, P., and J. Cote, 2013 Precise deposition of histone H2A.Z in chromatin for genome expression and maintenance. Biochim Biophys Acta 1819: 290–302.

5. Biswas, D., R. Dutta-Biswas, D. Mitra, Y. Shibata, B. D. Strahl et al., 2006 Opposing roles for Set2 and yFACT in regulating TBP binding at promoters. EMBO Journal 25: 4479–4489.

6. Bondarenko, V. A., L. M. Steele, A. Ujvari, D. A. Gaykalova, O. I. Kulaeva et al., 2006 Nucleosomes can form a polar barrier to transcript elongation by RNA polymerase II. Mol Cell 24: 469–479.

7. Briggs, S. D., M. Bryk, B. D. Strahl, W. L. Cheung, J. K. Davie et al., 2001 Histone H3 lysine 4 methylation is mediated by Set1 and required for cell growth and rDNA silencing in Saccharomyces cerevisiae. Genes & Development 15: 3286–3295.

8. Brunelle, M., C. Coulombe, C. Poitras, M. A. Robert, A. N. Markovits et al., 2015 Aggregate and Heatmap Representations of Genome-Wide Localization Data Using VAP, a Versatile Aggregate Profiler. Methods Mol Biol 1334: 273–298.

9. Burugula, B. B., C. Jeronimo, R. Pathak, J. W. Jones, F. Robert et al., 2014 Histone deacetylases and phosphorylated polymerase II C-terminal domain recruit Spt6 for cotranscriptional histone reassembly. Mol Cell Biol 34: 4115–4129.

10. Cheung, V., G. Chua, N. N. Batada, C. R. Landry, S. W. Michnick et al., 2008 Chromatin- and Transcription-Related Factors Repress Transcription from within Coding Regions throughout the Saccharomyces cerevisiae Genome. PLoS Biol 6: e277.

11. Close, D., S. J. Johnson, M. A. Sdano, S. M. McDonald, H. Robinson et al., 2011 Crystal structures of the S. cerevisiae Spt6 core and C-terminal tandem SH2 domain. J Mol Biol 408: 697–713.

12. Danko, C. G., N. Hah, X. Luo, A. L. Martins, L. Core et al., 2013 Signaling pathways differentially affect RNA polymerase II initiation, pausing, and elongation rate in cells. Mol Cell 50: 212–222.

13. Dechassa, M. L., A. Sabri, S. Pondugula, S. R. Kassabov, N. Chatterjee et al., 2010 SWI/SNF has intrinsic nucleosome disassembly activity that is dependent on adjacent nucleosomes. Mol Cell 38: 590–602.

14. Degennaro, C. M., B. H. Alver, S. Marguerat, E. Stepanova, C. P. Davis et al., 2013 Spt6 regulates intragenic and antisense transcription, nucleosome positioning, and histone modifications genome-wide in fission yeast. Mol Cell Biol 33: 4779–4792.

15. Dengl, S., A. Mayer, M. Sun and P. Cramer, 2009 Structure and in vivo requirement of the yeast Spt6 SH2 domain. J Mol Biol 389: 211–225.

16. Dion, M. F., T. Kaplan, M. Kim, S. Buratowski, N. Friedman et al., 2007 Dynamics of Replication-Independent Histone Turnover in Budding Yeast. Science 315: 1405–1408.

17. Duina, A. A., A. Rufiange, J. Bracey, J. Hall, A. Nourani et al., 2007 Evidence that the Localization of the Elongation Factor Spt16 Across Transcribed Genes Is Dependent Upon Histone H3 Integrity in Saccharomyces cerevisiae. Genetics 177: 101–112.

18. Dutta, A., M. Gogol, J.-H. Kim, M. Smolle, S. Venkatesh et al., 2014 Swi/Snf dynamics on stress-responsive genes is governed by competitive bromodomain interactions. Genes & Development 28: 2314–2330.

19. Endoh, M., W. Zhu, J. Hasegawa, H. Watanabe, D. K. Kim et al., 2004 Human Spt6 stimulates transcription elongation by RNA polymerase II in vitro. Mol Cell Biol 24: 3324–3336.

20. Fleming, A. B., C.-F. Kao, C. Hillyer, M. Pikaart and M. A. Osley, 2008 H2B Ubiquitylation Plays a Role in Nucleosome Dynamics during Transcription Elongation. Molecular Cell 31: 57–66.

21. Formosa, T., 2012 The role of FACT in making and breaking nucleosomes. Biochim Biophys Acta 1819: 247–255.

22. Formosa, T., P. Eriksson, J. Wittmeyer, J. Ginn, Y. Yu et al., 2001 Spt16-Pob3 and the HMG protein Nhp6 combine to form the nucleosome-binding factor SPN. Embo j 20: 3506–3517.

23. Ginsburg, D. S., C. K. Govind and A. G. Hinnebusch, 2009 NuA4 Lysine Acetyltransferase Esa1 Is Targeted to Coding Regions and Stimulates Transcription Elongation with Gcn5. Mol. Cell. Biol. 29: 6473–6487.

24. Govind, C. K., D. Ginsburg and A. G. Hinnebusch, 2012 Measuring Dynamic Changes in Histone Modifications and Nucleosome Density during Activated Transcription in Budding Yeast. Methods Mol Biol 833: 15–27.

25. Govind, C. K., H. Qiu, D. S. Ginsburg, C. Ruan, K. Hofmeyer et al., 2010 Phosphorylated Pol II CTD recruits multiple HDACs, including Rpd3C(S), for methylation-dependent deacetylation of ORF nucleosomes. Mol Cell 39: 234–246.

26. Govind, C. K., F. Zhang, H. Qiu, K. Hofmeyer and A. G. Hinnebusch, 2007 Gcn5 promotes acetylation, eviction, and methylation of nucleosomes in transcribed coding regions. Mol Cell 25: 31–42.

27. Gurard-Levin, Z. A., J. P. Quivy and G. Almouzni, 2014 Histone chaperones: assisting histone traffic and nucleosome dynamics. Annu Rev Biochem 83: 487–517.

28. Hassan, A. H., K. E. Neely and J. L. Workman, 2001 Histone acetyltransferase complexes stabilize swi/snf binding to promoter nucleosomes. Cell 104: 817–827.

29. Hondele, M., and A. G. Ladurner, 2011 The chaperone-histone partnership: for the greater good of histone traffic and chromatin plasticity. Curr Opin Struct Biol 21: 698–708.

30. Hondele, M., T. Stuwe, M. Hassler, F. Halbach, A. Bowman et al., 2013 Structural basis of histone H2A-H2B recognition by the essential chaperone FACT. Nature 499: 111–114.

31. Hsieh, F. K., O. I. Kulaeva, S. S. Patel, P. N. Dyer, K. Luger et al., 2013 Histone chaperone FACT action during transcription through chromatin by RNA polymerase II. Proc Natl Acad Sci U S A 110: 7654–7659.

32. Hughes, T. R., M. J. Marton, A. R. Jones, C. J. Roberts, R. Stoughton et al., 2000 Functional discovery via a compendium of expression profiles. Cell 102: 109–126.

33. Ivanovska, I., P. E. Jacques, O. J. Rando, F. Robert and F. Winston, 2011 Control of chromatin structure by spt6: different consequences in coding and regulatory regions. Mol Cell Biol 31: 531–541.

34. Jamai, A., A. Puglisi and M. Strubin, 2009 Histone Chaperone Spt16 Promotes Redeposition of the Original H3-H4 Histones Evicted by Elongating RNA Polymerase. Molecular Cell 35: 377–383.

35. Jeronimo, C., S. Watanabe, C. D. Kaplan, C. L. Peterson and F. Robert, 2015 The Histone Chaperones FACT and Spt6 Restrict H2A.Z from Intragenic Locations. Mol Cell 58: 1113–1123.

36. Jimeno-Gonzalez, S., F. Gomez-Herreros, P. M. Alepuz and S. Chavez, 2006 A gene-specific requirement for FACT during transcription is related to the chromatin organization of the transcribed region. Mol Cell Biol 26: 8710–8721.

37. Kaplan, C. D., M. J. Holland and F. Winston, 2005 Interaction between transcription elongation factors and mRNA 3’-end formation at the Saccharomyces cerevisiae GAL10-GAL7 locus. J Biol Chem 280: 913–922.

38. Kaplan, C. D., L. Laprade and F. Winston, 2003 Transcription elongation factors repress transcription initiation from cryptic sites. Science 301: 1096–1099.

39. Kaplan, C. D., J. R. Morris, C. Wu and F. Winston, 2000 Spt5 and spt6 are associated with active transcription and have characteristics of general elongation factors in D. melanogaster. Genes Dev 14: 2623–2634.

40. Kato, H., K. Okazaki, T. Iida, J. Nakayama, Y. Murakami et al., 2013 Spt6 prevents transcription-coupled loss of posttranslationally modified histone H3. Sci Rep 3.

41. Kemble, D. J., L. L. McCullough, F. G. Whitby, T. Formosa and C. P. Hill, 2015 FACT Disrupts Nucleosome Structure by Binding H2A-H2B with Conserved Peptide Motifs. Mol Cell 60: 294–306.

42. Kireeva, M. L., W. Walter, V. Tchernajenko, V. Bondarenko, M. Kashlev et al., 2002 Nucleosome remodeling induced by RNA polymerase II: loss of the H2A/H2B dimer during transcription. Mol Cell 9: 541–552.

43. Krogan, N. J., M. Kim, S. H. Ahn, G. Zhong, M. S. Kobor et al., 2002 RNA polymerase II elongation factors of Saccharomyces cerevisiae: a targeted proteomics approach. Mol Cell Biol 22: 6979–6992.

44. Lee, C.-K., Y. Shibata, B. Rao, B. D. Strahl and J. D. Lieb, 2004 Evidence for nucleosome depletion at active regulatory regions genome-wide. Nat Genet 36: 900–905.

45. Li, B., M. Carey and J. L. Workman, 2007 The role of chromatin during transcription. Cell 128: 707–719.

46. Liu, J., J. Zhang, Q. Gong, P. Xiong, H. Huang et al., 2011 Solution structure of tandem SH2 domains from Spt6 protein and their binding to the phosphorylated RNA polymerase II C-terminal domain. J Biol Chem 286: 29218–29226.

47. Mason, P. B., and K. Struhl, 2003 The FACT complex travels with elongating RNA polymerase II and is important for the fidelity of transcriptional initiation in vivo. Mol Cell Biol 23: 8323–8333.

48. Mayer, A., M. Heidemann, M. Lidschreiber, A. Schreieck, M. Sun et al., 2012 CTD tyrosine phosphorylation impairs termination factor recruitment to RNA polymerase II. Science 336: 1723–1725.

49. Mayer, A., M. Lidschreiber, M. Siebert, K. Leike, J. Soding et al., 2010 Uniform transitions of the general RNA polymerase II transcription complex. Nat Struct Mol Biol 17: 1272–1278.

50. McCullough, L., Z. Connell, C. Petersen and T. Formosa, 2015 The Abundant Histone Chaperones Spt6 and FACT Collaborate to Assemble, Inspect, and Maintain Chromatin Structure in Saccharomyces cerevisiae. Genetics 201: 1031–1045.

51. Mnaimneh, S., A. P. Davierwala, J. Haynes, J. Moffat, W. T. Peng et al., 2004 Exploration of essential gene functions via titratable promoter alleles. Cell 118: 31–44.

52. Nguyen, H. T., W. Wharton, 2nd, J. A. Harper, J. R. Dornhoffer and A. A. Duina, 2013 A nucleosomal region important for ensuring proper interactions between the transcription elongation factor Spt16 and transcribed genes in Saccharomyces cerevisiae. G3 (Bethesda) 3: 929–940.

53. Pavri, R., B. Zhu, G. Li, P. Trojer, S. Mandal et al., 2006 Histone H2B monoubiquitination functions cooperatively with FACT to regulate elongation by RNA polymerase II. Cell 125: 703–717.

54. Perales, R., B. Erickson, L. Zhang, H. Kim, E. Valiquett et al., 2013 Gene promoters dictate histone occupancy within genes. EMBO J 32: 2645–2656.

55. Prendergast, J. A., L. E. Murray, A. Rowley, D. R. Carruthers, R. A. Singer et al., 1990 Size selection identifies new genes that regulate Saccharomyces cerevisiae cell proliferation. Genetics 124: 81–90.

56. Ransom, M., S. K. Williams, M. L. Dechassa, C. Das, J. Linger et al., 2009 FACT and the proteasome promote promoter chromatin disassembly and transcriptional initiation. J Biol Chem 284: 23461–23471.

57. Reinke, H., P. D. Gregory and W. Horz, 2001 A Transient Histone Hyperacetylation Signal Marks Nucleosomes for Remodeling at the *PHO8* Promoter in Vivo. Mol.Cell. 7: 529–538.

58. Simic, R., D. L. Lindstrom, H. G. Tran, K. L. Roinick, P. J. Costa et al., 2003 Chromatin remodeling protein Chd1 interacts with transcription elongation factors and localizes to transcribed genes. Embo J 22: 1846–1856.

59. Spain, M. M., S. A. Ansari, R. Pathak, M. J. Palumbo, R. H. Morse et al., 2014 The RSC complex localizes to coding sequences to regulate Pol II and histone occupancy. Mol Cell 56: 653–666.

60. Squazzo, S. L., P. J. Costa, D. L. Lindstrom, K. E. Kumer, R. Simic et al., 2002 The Paf1 complex physically and functionally associates with transcription elongation factors in vivo. EMBO J 21: 1764–1774.

61. Strahl, B. D., P. A. Grant, S. D. Briggs, Z. W. Sun, J. R. Bone et al., 2002 Set2 is a nucleosomal histone H3-selective methyltransferase that mediates transcriptional repression. Mol Cell Biol 22: 1298–1306.

62. Stuwe, T., M. Hothorn, E. Lejeune, V. Rybin, M. Bortfeld et al., 2008 The FACT Spt16 “peptidase” domain is a histone H3-H4 binding module. Proc Natl Acad Sci U S A 105: 8884–8889.

63. Takahata, S., Y. Yu and D. J. Stillman, 2009 FACT and Asf1 Regulate Nucleosome Dynamics and Coactivator Binding at the HO Promoter. Molecular Cell 34: 405–415.

64. van Bakel, H., K. Tsui, M. Gebbia, S. Mnaimneh, T. R. Hughes et al., 2013 A Compendium of Nucleosome and Transcript Profiles Reveals Determinants of Chromatin Architecture and Transcription. PLoS Genet 9: e1003479.

65. VanDemark, A. P., H. Xin, L. McCullough, R. Rawlins, S. Bentley et al., 2008 Structural and functional analysis of the Spt16p N-terminal domain reveals overlapping roles of yFACT subunits. J Biol Chem 283: 5058–5068.

66. Venkatesh, S., M. Smolle, H. Li, M. M. Gogol, M. Saint et al., 2012 Set2 methylation of histone H3 lysine 36 suppresses histone exchange on transcribed genes. Nature 489: 452–455.

67. Winkler, D. D., U. M. Muthurajan, A. R. Hieb and K. Luger, 2011 Histone chaperone FACT coordinates nucleosome interaction through multiple synergistic binding events. J Biol Chem 286: 41883–41892.

68. Xin, H., S. Takahata, M. Blanksma, L. McCullough, D. J. Stillman et al., 2009 yFACT Induces Global Accessibility of Nucleosomal DNA without H2A-H2B Displacement. Molecular Cell 35: 365–376.

